# Novel Neuromuscular Controllers with Simplified Muscle Model and Enhanced Reflex Modulation: A Comparative Study in Hip Exoskeletons

**DOI:** 10.1101/2024.05.10.593488

**Authors:** Ali Reza Manzoori, Sara Messara, Andrea Di Russo, Auke Ijspeert, Mohamed Bouri

## Abstract

Neuromuscular controllers (NMCs) offer a promising approach to adaptive and task-invariant control of exoskeletons for walking assistance, leveraging the bioinspired models based on the peripheral nervous system. This article expands on our previous development of a novel structure for NMCs with modifications to the virtual muscle model and reflex modulation strategy. The modifications consist firstly of simplifications to the Hill-type virtual muscle model, resulting in a more straightforward formulation and reduced number of parameters; and secondly, using a finer division of gait subphases in the reflex modulation state machine, allowing for a higher degree of control over the shape of the assistive profile. Based on the proposed general structure, we present two controller variants for hip exoskeletons, with four- and five-state reflex modulations (NMC-4 and NMC-5). We used an iterative data-driven approach with two tuning stages (i.e., muscle parameters and reflex gains) to determine the controller parameters. Biological joint torque profiles and optimal torque profiles for metabolic cost reduction were used as references for the final tuning outcome. Experimental testing under various walking conditions demonstrated the capability of both variants for adapting to the locomotion task with minimal parameter adjustments, mostly in terms of timing. Furthermore, NMC-5 exhibited better alignment with biological and optimized torque profiles in terms of timing characteristics and relative magnitudes, resulting in less negative mechanical work. These findings firstly validate the adequacy of the simplified muscle model for assistive controllers, and demonstrate the utility of a more nuanced reflex modulation in improving the assistance quality.

## 1 Introduction

Lower-limb exoskeletons are increasingly gaining traction for applications outside of research labs. In addition to use in rehabilitation clinics (Plaza et al., 2023) and facilitating functional mobility for individuals with severe impairments (van Dijsseldonk et al., 2020), exoskeletons are being considered for assisting workers in industrial (Kim et al., 2019) and construction sites (de Looze et al., 2016), improving athletic performance (Moon et al., 2023), and even maintaining fitness in low-gravity environments (Rea et al., 2013). Although specific performance requirements vary across these different use cases, a common challenge is effectively bridging the gap between human intent and the actions of the assistive device (Lobo-Prat et al., 2014). While many of the existing exoskeleton control strategies can effectively provide assistance under controlled settings, they often struggle in adapting to the diverse needs and dynamic contexts of real-world scenarios, due to the reliance on predefined parameters or assistance patterns. This restricts the exoskeleton’s ability to seamlessly adjust its support to different locomotion tasks (e.g., walking, stair climbing, sit-to-stand) and individual user characteristics (e.g., strength, gait patterns).

One way to overcome this limitation is adding a higher-level layer to adjust the controller based on recognition of the terrain and its features (Al-dabbagh and Ronsse, 2020), detection of the locomotion task (Moreira et al., 2022) and its characteristics (Slade et al., 2022; Medrano et al., 2023), or measurements of the users’ response to variations in assistance (human-in-the-loop optimisation approaches) (Ding et al., 2018; Song and Collins, 2021). While these approaches have shown promising outcomes, they come with trade-offs. The added complexity demands more sensory information, higher computational loads, and extensive data and training.

Another approach is to rely on control methods that inherently adapt to the user and task, without the need for explicit detection and parameter adjustments. That is, rather than using predefined assistance patterns or trajectories, the assistive torque/force emerges in real-time from the interaction of the device with the user. A basic example of such methods which was proposed early in the exoskeleton literature is proportional myoelectric control (Ferris et al., 2005). In this method, the exoskeleton torque is directly determined by the muscle activities of the user. Despite the straightforward concept and positive experimental outcomes, this method is hindered by issues associated with EMG sensing, such as sensitivity to electrode placement, susceptibility to noise and variability, and the necessity for user-specific calibration (Türker, 1993; Cimolato et al., 2022). Model-based methods such as “integral admittance shaping” (Nagarajan et al., 2016) and “energy shaping” (Lin et al., 2022) leverage dynamical models of walking mechanics. By manipulating the general dynamics of the human-exoskeleton system, these methods can provide task-agnostic assistance. However, implementation of such methods is challenging in practice, mainly due to their dependence on accurate models and parameters such as segment inertias, which are difficult to obtain accurately. Alternatively, there are heuristic methods of mapping sensory information to assistive action such as the one proposed by Lim et al. (2019) for hip exoskeletons. While these methods tend to be more practically feasible without the need for complex models or sensors, the lack of a systematic approach limits them to a specific joint and offers a narrower scope for fine-tuning the provided assistance.

Inspired by the remarkable versatility of human locomotion, exoskeleton control strategies based on neuromuscular and biomechanical models of human gait provide a different avenue for improving controller adaptability (Firouzi et al., 2023). A main group of these methods was initiated by the early work of Geyer et al. (2003); Geyer and Herr (2010) in which a simplified body model actuated by virtual muscles, driven by local feedback loops mimicking reflexes of the peripheral nervous system, generated stable and robust human-like gaits in simulation. Controllers based on this idea, also known as neuromuscular controllers (NMCs) were implemented on prostheses (Eilenberg et al., 2010) and later on exoskeletons (Dzeladini et al., 2016), showing inherent adaptability without the need for extensive sensory inputs (Tagliamonte et al., 2022). Thanks to the modularity of the model, it has been extended to assist different joints and degrees of freedom, including sagittal hip-knee (Wu et al., 2017) and ankle (Tamburella et al., 2020; Shafer et al., 2021), and Meijneke et al. (2021) have extended it by including abduction/adduction assistance too. In a recent study (Afschrift et al., 2023), the controller has been augmented with a reflex driven by the center of mass velocity deviations from a standard trajectory in addition to local reflexes, and implemented on an ankle exoskeleton to also assist balance recovery after perturbations. A similar implementation with center of mass reflexes for standing balance assistance has also been reported (Yin et al., 2022).

Despite differences in reflex mappings and configuration of the joints, all of the mentioned implementations of NMC use a two-phase (i.e., stance/swing) reflex modulation and a Hill-type (Miller, 2018) muscle model, similar to the original structure introduced by Geyer and Herr (2010). However, some studies exploring neuromechanical models in simulation have utilized more nuanced reflex modulations based on subphases within stance and swing for finer control of generated behaviors (Wang et al., 2012; Ong et al., 2019). Furthermore, the Hill-type model was originally designed to replicate the behavior of biological muscles. While achieving high fidelity with respect to biological systems can be beneficial for predictive and explanatory objectives in simulation studies, the added complexity might not be warranted in practical applications for controlling assistive devices. In addition to the added computational complexity and the need for numerical solutions, the Hill model can also cause numerical issues (Van der Noot et al., 2014; Yeo et al., 2023).

Based on these observations, we proposed modifications to the commonly used NMC structure, simplifying the virtual muscle model and using a more fine-grained reflex modulation (Messara et al., 2023). The simplification of the muscle model aims to reduce the computational complexity without loss of functionality. In contrast, the added nuance to the reflex modulation structure is intended to improve the adaptability of the controller’s output and facilitate tailoring the torque profiles to specific needs. In this article, we give a more thorough description of the novel structure, and detail our parameter tuning strategy. We also test and compare the performance of two variants of the controller designed for hip exoskeletons, with different numbers of phases for reflex modulation to evaluate the utility of using more phases. A more in-depth analysis of the results of assisted walking experiments is also given.

## 2 Materials and Methods

### 2.1 Controller Structure And Modifications

The general structure of NMCs (see Figure 1 for an example) can be divided into two parts: the high-level “neural” layer, where the feedback mappings (termed “reflexes”) between sensory inputs and control actions are defined; and the low-level “muscular” layer, where the action signals from the neural layer are translated into joint torque commands using virtual muscles. Our proposed modifications to the NMC structure involve both layers, and will be described separately here. For each layer, we will provide the general formulation, and then describe our proposed modifications. We will use the same notations as Geyer and Herr (2010) unless stated otherwise.

**Figure 1:**
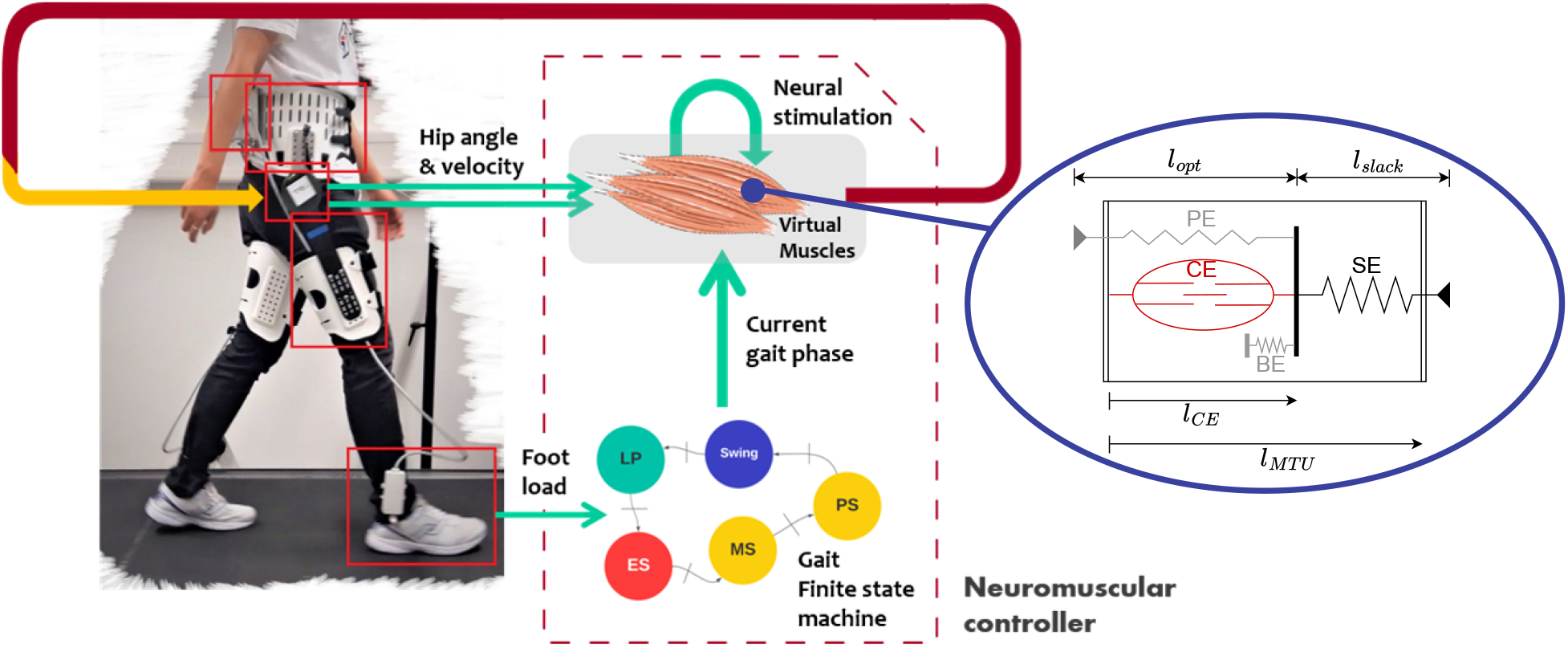
The schematic structure of the NMC (left) and the virtual muscle model (right).

#### 2.1.1 Virtual Muscles

The Hill-type muscle models are a class of phenomenological models of the biological muscle initiated by the early work of Hill (1938), which have been widely used in computational models of animal movement. These models represent the muscle-tendon unit (MTU) with a combination of active and passive nonlinear elastic elements (Miller, 2018). The commonly used configuration in NMCs (the one used by Geyer and Herr (2010)) consists of four elastic elements (Figure 1, right). Three of the elements are passive; namely, the serial element (SE), the parallel element (PE) and the buffer element (BE). PE and BE only act when the MTU length is outside of its normal operating range. The last element, termed the contractile element (CE), is the only active element which generates muscle contractions. The force generated by the passive (NLP) elements at the contraction is given by the force-length relationship (Geyer and Herr, 2010):

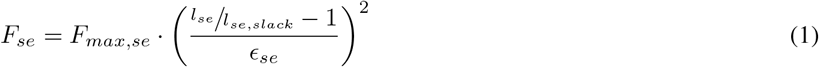

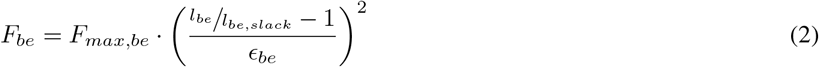

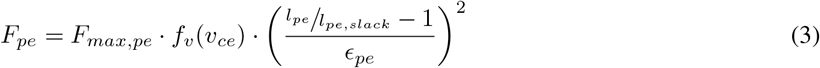

Where *l*_*nlp*_^3^ is the length of the NLP element, *l*_*nlp,slack*_^4^ is its slacking length, *ϵ*_*nlp*_^5^ the reference strain and *F*_*max,nlp*_ the maximum force.

The CE force at the contraction, denoted as *F*_*ce*_, is calculated based on the muscle activation *Act*, the force-length relationship *f*_*l*_ and the force-velocity relationship *f*_*v*_, described as:

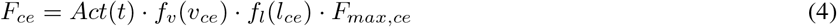

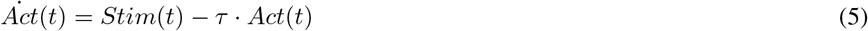

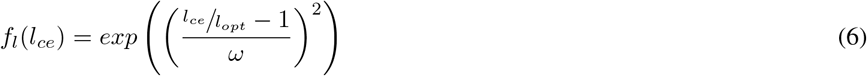

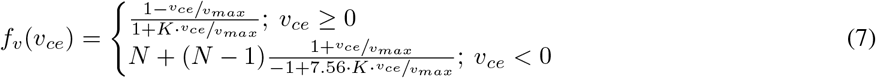

The equations involve the two internal state variables of the CE, namely, the CE length (*l*_*ce*_) and velocity (*v*_*ce*_), in addition to the activation (*Act*) signal, which is determined by the stimulation (*Stim*) signal (received from the neural layer) through the dynamical equation 5 with the time constant *τ* . In the force-length relationship (Equation 6), the constant *ω* determines the width of the Gaussian. In the force-velocity relationship (Equation 7), *K* is a curvature constant and *N* is the normalized MTU force, *F*_ce_*/F*_max,ce_. Finally, the generated force at each MTU is given by:

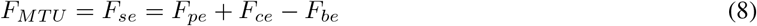

The third part of the model establishes the links between the mechanical state of the joint and that of each connected MTU. These relationships are specific to each joint and deduced from the geometry of the joint model. For the hip joint:

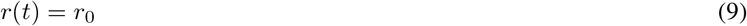

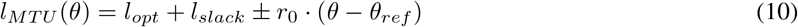

The choice of the operation for the *±* depends on the type of the muscle: + for the flexor and *−* for the extensor. The equations for the knee and ankle muscles (in the case of bi-articular muscles whose states present a coupling of the two joints) are:

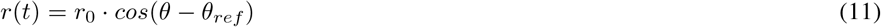

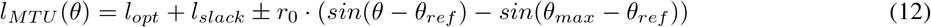

Again, the *±* sign depends on the type of the muscle, but for this group of muscles *−* corresponds to flexor and + to extensor muscles.

The final torque around the joints is finally given by:

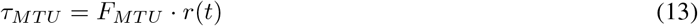

Finding the MTU force requires resolving six state variables: one neural state variable (CE activation) which depends directly on the neural stimulation, and five mechanical state variables, namely the lengths of the BE, SE, PE and CE elements and the velocity of the CE element. Solving for the mechanical state variables is more challenging as the number of the mechanical equations (three) is less than the number of the unknown variables (five). The challenge is further aggravated by the nonlinear and highly coupled nature of the system.

Geyer and Herr (2010) adopted an iterative approach that uses the precedent value of the CE length to compute all the forces and then invert the *f*_*v*_ relationship to find *v*_*ce*_. The next value of *l*_*ce*_ is then obtained by numerically integrating *v*_*ce*_. As this method uses an iterative pattern with inversions and numerical integration, this makes it less accurate and less adapted to real-time calculations. For these reasons, we propose two simplifications to the model that lead to a more efficient computational approach.

##### Simplifications of the model

In order to solve the mechanical states algebraically, we assumed the SE element (which roughly captures the series elasticity of the tendon) to have a constant length. This is motivated by the high biological stiffness of the tendon that is reported in the literature to be around 150 kN *·* m^*−*1^ (Maganaris and Paul, 1999). By making this assumption, the internal degrees of freedom are reduced, which allows to exploit the analytical derivative of the CE length to obtain its velocity:

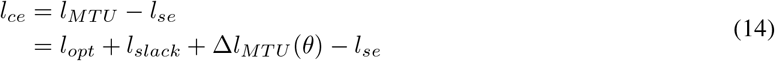

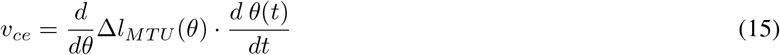

The expression of Δ*l*_*MTU*_ (*θ*) depends on the considered joint, according to Equations 10 and 12. For example, in the case of the hip joint, from Equation 10:

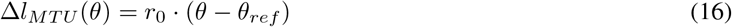

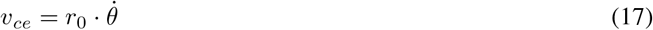

We call this computational strategy the derivative approach, which contrasts with the integral approach for obtaining *l*_*ce*_ from *v*_*ce*_, as proposed by Geyer and Herr (2010) and utilized in the classic NMC structure.

The force generated at the joint level is approximated as:

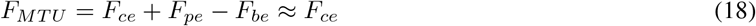

The last approximation comes from the length equality between the BE and PE elements, which makes the subtraction of their respective forces negligible. Furthermore, these two elements only act when the MTU is over-stretched or over-slacked, and thus do not have a major effect on the generated forces.

##### Choice of virtual muscles and joint torque

In this study, we target a hip exoskeleton and therefore only the hip joint is considered in the neuromuscular model. Around the hip joint, the model of Geyer and Herr (2010) considered two extensors, namely, hamstring (HAMS) and gluteus maximus (GLU), in addition to the hip flexor group (HFL). The same muscles were used by Ong et al. (2019), but the HFL was decomposed into rectus femoris (RF) and iliopsoas (ILPS). Since the exoskeleton used in this study provides actuation and sensing only at the hip level, we only considered the mono-articular muscles solely affecting the hip, i.e., GLU as the extensor and ILPS as the flexor, to avoid dependence on the state of the knee joint.

During tuning and pilot testing, we observed that without re-tuning of the parameters, the amplitude of the generated assistive torques did not sufficiently adapt to various walking conditions. Therefore, we introduced a scaling gain *G*_*s,v*_, which was manually tuned according to the walking speed and ground inclination. The total applied torque is thus given by:

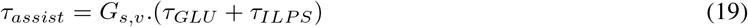

A detailed block diagram representation of the controller is shown in Supplementary Figure 1. For the experiments, the output of this equation was sent through a low-pass filter (first-order, *f*_*c*_ = 20 Hz) in order to enhance the smoothness of the torque profile and prevent discomfort to the user.

#### 2.1.2 Reflex modulation and state machine

The stimulation signal used in Equation 5 is determined in the neural layer, based on one or a combination of reflexes for each muscle, with this general expression:

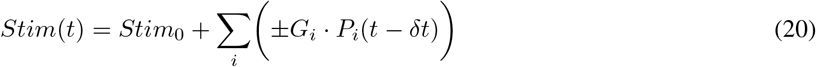

Where *Stim*_0_ is a basal stimulation, *G*_*i*_’s are reflex gains, and *P*_*i*_’s are reflex sensory inputs delayed by a signal propagation time *δt*. Typically used reflex sensory inputs are muscle forces, lengths and rates of lengthening, but constant values have also been used. In this work, we chose to rely only on length feedback and constant inhibition. This choice is motivated mostly by the smoothness of the length signal. Moreover, it has been shown that length and velocity feedback were sufficient to adapt to different walking speeds (Ong et al., 2019). The general expression of *Stim* in our case thus becomes:

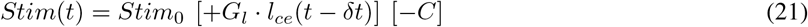

where *G*_*l*_ is the length reflex gain, and *C* is a constant inhibition term. For the hip muscles, we used *δt* = 5 ms is a constant simulating the neural signal delays, and *C* is a constant inhibition term.

The neural stimulation of the leg muscles during the gait cycle exhibits trends that are dependent on the gait phase (Nielsen and Sinkjær, 2002). In accordance with this observation, the goal of reflex modulation is to determine the reflexes included in the summation term on the right hand side of Equation 20 in each subphase of gait. For this reason, a finite state machine (FSM) composed of gait phases is a good candidate for reflex modulation. In addition, FSMs with state transitions triggered by gait events do not require perfect periodicity or prior knowledge of the walking speed or cadence, unlike time-based synchronisation methods. In their initial study, Geyer and Herr (2010) considered only two states, corresponding to the stance and swing phases, which is a rather coarse decomposition of the gait cycle.

In our implementation, we opted for a more detailed state decomposition that could better encapsulate the continuous trends of stimulation during gait. The design of the FSM and its state transition criteria must account for variations in gait as a result of environmental factors, as well as the inter-person variability in gait patterns. Hence, in this work we rely on robustly detectable events with little inter-person variability based on ground contact status and zero crossings of the angular velocity. We propose two variants of the controller with four and five states, named NMC-4 and NMC-5 respectively, as detailed in what follows.

##### Four-state FSM (NMC-4)

This state machine includes the following states: Early Stance (ES), Mid-stance and Pre-swing (MS-PS), Swing (S) and landing preparation (LP). The events for detecting the state transitions are the ipsilateral heel-strike (LP to ES), the contralateral toe-off (ES to MS-PS), the ilpsilateral toe-off (MS-PS to S), and the flexion-to-extension zero crossing of the ipsilateral hip angular velocity (S to LP).

**Table 1:** NMC-4 reflex modulations for each FSM state, displaying the types of reflexes (positive length, *L*^+^, or constant inhibition, *C*^*−*^) along with the numerical values (in brackets).

| Phase<br>Muscle | ES | MS – PS | S | LP |
| --- | --- | --- | --- | --- |
| GLU | $L^+[0.8]$ | $L^+[0.8], C^-[0.08]$ | $basal[0.01]$ | $L^+[0.8]$ |
| ILPS | $basal[0.01]$ | $L^+[0.6]$ | $L^+[0.6]$ | $basal[0.01]$ |

##### Five-state FSM (NMC-5)

The considered states in this FSM are: Early Stance (ES), Mid-stance (MS), Pre-swing (PS), Swing (S) and landing preparation (LP). The events are similar to those of the four-state FSM, in addition to the contra-lateral heel-strike for the transition from MS to PS.

**Table 2:**
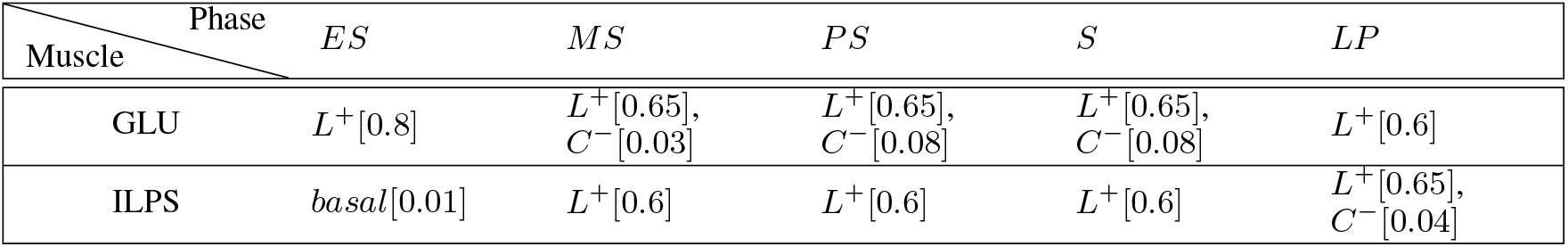
NMC-5 reflex modulations for each FSM state, displaying the types of reflexes (positive length, *L*^+^, or constant inhibition, *C*^*−*^) along with the numerical values (in brackets).

### 2.2 Tuning Procedures

The muscular model consists of eighteen parameters per virtual muscle, plus six reflex gains per leg for the NMC-4 variant and thirteen gains per leg for NMC-5. Here, we describe the detailed tuning procedure, a summary of which was provided in our previous work (Messara et al., 2023).

#### 2.2.1 Virtual muscle parameter tuning

For the tuning of the virtual muscle parameters, we used data-driven simulations, in which prerecorded gait data was provided as input to the controller in a simulated environment. We relied on two former studies for the initial values; the one of Geyer and Herr (2010) in which the gains were manually tuned in simulation, and the study of Ong et al. (2019) which optimised a multi-objective function including the gross cost of transport, falling and ligament injuries minimisation and head tilting and step speeds stability maximisation. We followed an iterative approach, consisting in analysing the resulting torques and the various state variables (*l*_*ce*_, *v*_*ce*_, *τ*_*MTU*_, *Stim, Act*, etc.). Two torque profiles were used as the benchmarks for this comparison; i.e., the estimated torque profiles of the human hip joint (Winter, 1991; Reznick et al., 2021), to which we will refer as “biological profiles”, and those obtained in human in the loop (HitL) assistance torque optimisation studies (Ding et al., 2018; Franks et al., 2022), which we will address as “HitL profiles”. This approach was used to isolate a set of focal parameters to be re-tuned, varying one parameter per iteration. The identified focal parameters and their effects were:

- the optimal CE length (*l*_*opt*_) in Equation 6, which affects the Gaussian peak of the force-length relationship, and the reference angle (*θ*_*ref*_) which influences the MTU force lever. These two parameters correlate to the timing of the flexion and extension torque peaks and are thus crucial in setting the torque profile;
- the maximum speed (*v*_*max*_) and the constant *K* in Equation 7, which shape the slope of the force-velocity relationship and control accordingly the sensitivity of the torque profile to velocity variations;
- the maximum force (*F*_*max,ce*_) in Equation 4, which sets the magnitudes of the muscular force and joint torque. Since only two muscles are considered in the model, it was not possible to keep the biological force ratio between the GLU and the ILPS as this would have led to a disequilibrium between the flexion and the extension torques. Thus, the maximum forces ratio was set equal in order to compensate the absence of the other muscles.
- the cut-off period (*τ*) in Equation 5 defines the attenuation between the muscle stimulation and activation and accordingly the torque profile smoothing;
- the choice of SE length (*l*_*se*_) was tuned to a mean working value since it is assumed constant in the simplified model.

The remaining model parameters had minor adjustments to their initial values. The numerical values of the parameters are reported in Supplementary Table 1.

#### 2.2.2 Neural reflex gain tuning

Tuning the reflex gains was more challenging, given that the chosen types and combination of reflexes were different from previous studies. In addition, the gains highly influence the magnitude of the generated torques, and thus needed to be adapted to the hardware capacities of the target exoskeleton in terms of peak and nominal torques of the actuators. We used a two-stage simulation with hand tuning. First, we performed hardware in the loop simulation which consisted in playing pre-recorded gait kinematic profiles on the exoskeleton’s embedded computer. Second, system in the loop with the exoskeleton was used; the torques were simulated on the exoskeleton in real time based on the kinematics measured online, but it was not applied yet on the motors. The generated torque was monitored using a desktop application connected to the embedded computer, then MATLAB was used for further analysis. The tuning criteria of the inhibition constants and gains were, first, the continuity and smoothness of the torque profile despite the discrete state transitions and second, the mimicry of the biological and HitL torque profile in terms of curve tendency, characteristic times and magnitude ratio (Winter, 1991; Franks et al., 2021). Regarding the slope (s) and velocity (v) adaptation gain (*G*_*s,v*_), we used the average ratio between the biological torque profiles of different inclines and speed groups, compared to a baseline walking condition (walking speed of 1.25 m *·* s^*−*1^ and 0% slope) (Reznick et al., 2021; Winter, 1991). The values of *G*_*s,v*_ were 0.8, 1, 1.15 for the slow, normal and high speed groups, respectively and 1.15 for the incline (the same gain was applied for all inclines not exceeding 15%).

### 2.3 Experimental setup and protocol

Nine healthy participants (5 females, 4 males, age: 27 *±* 2.3 years, weight: 65.0 *±* 8.4 kg, height: 171.8 *±* 4.7 cm) were recruited for the experimental testing. The protocol was reviewed and approved by the EPFL Human Research Ethics Committee. The participants provided their written informed consent prior to partaking in the experiment.

An autonomous hip exoskeleton (e-Walk V1) was used as the implementation platform for the controllers. The exoskeleton weighs 5 kg and can deliver torques of up to 35 N *·* m around the frontal axis in active mode. In passive mode, the joints are easily backdrivable thanks to efficient planetary reducers with a 6:1 ratio. Flexibility of the thigh segments around the sagittal axis allows hip abduction in the limited range required for walking. Hip flexion/extension angles and angular velocities are measured using the encoders of the motors, and the output torques are estimated from motor currents. Additionally, foot contact status is determined using insole pressure sensors. All control and data logging operations were performed on the embedded computer of the exoskeleton at 500 Hz. More details about the device can be found in our previous article (Messara et al., 2023). A treadmill with adjustable speed and inclination (N-Mill, Forcelink B.V., Netherlands) was used for all of the experimental conditions. The experimental setup is schematically represented in Figure 2A.

**Figure 2:**
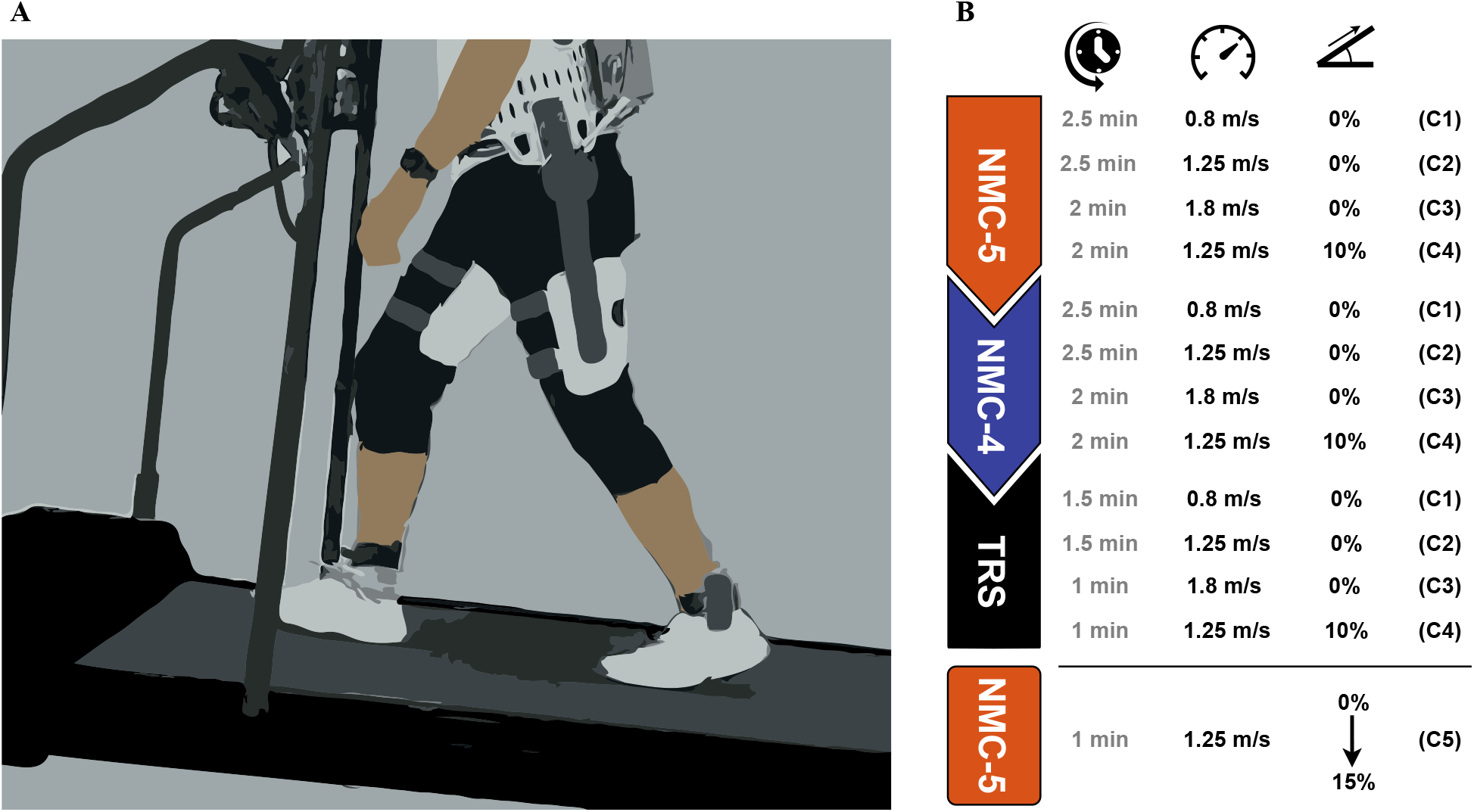
Experimental setup and procedure. **(A)** Schematic illustration of the experimental setup. **(B)** The order and specifications of the experimental conditions.

The experimental procedure consisted of walking on the treadmill at various speeds and inclinations while wearing the exoskeleton, starting with 0% inclination at 0.8 m *·* s^*−*1^ (C1), 1.25 m *·* s^*−*1^ (C2), and 1.8 m *·* s^*−*1^ (C3), and then 1.25 m *·* s^*−*1^ with 10% inclination (C4). We will refer to C1–C4 as the “constant” conditions, since the treadmill speed and inclination were constant in each one. The participants went through the constant conditions first with NMC-5, followed by NMC-4. The same conditions were also repeated with the exoskeleton in transparent (zero-torque) mode (TRS) to measure each participant’s baseline kinematics. The chronological order of the conditions was the same for all participants, as shown in Figure 2, but the durations were slightly shorter in the TRS set. Lastly, in order to test the adaptation of assistance to continuous changes in inclination, a final condition was tested during which the inclination of the treadmill gradually increased from 0 to 15% over 1 min (C5). We will refer to this condition as the “variable inclination” condition. While both variants were tested under this condition, we will only present the results for NMC-5 since it consistently showed a better performance in the constant conditions.

The only parameter that was modified between the conditions was the linear scaling gain *G*_*s,v*_, which was manually changed to improve the adaptation of assistance amplitude as explained in 2.2.2.

### 2.4 Data processing and analysis

All of the gathered data were first visually inspected for integrity prior to processing. The data for three participants (P3, P4 and P6) in the C5 condition were excluded from analysis due to issues in ground contact sensing. Signals from the insole pressure sensors were used for heel-strike detection and stride segmentation. To analyze the generated assistive torques, we directly used the torque commands calculated from Equation 19 (after low-pass filtering, as explained in Section 2.1.1). To calculate the mechanical power output of the exoskeleton, however, we used the measured torques (based on actuator currents). All torque and power values were normalized by the participants’ body mass. Due to the symmetry between left and right sides, we only used the data for the left leg in our analyses. Before calculating the average profiles, we removed the outlier strides, identified as those including samples that were more than three scaled median absolute deviations away from the local median. To obtain the means and standard deviations across all participants, we firstly calculated the mean values over all strides for each participant and then calculated the grand mean and standard deviation from the participant means. Angles and torques in the direction of extension were taken to be positive. All data processing was carried out using MATLAB.

## 3 Results

### 3.1 Assistive torques and powers

Overall, the generated assistive torques by both the NMC-4 and NMC-5 variants displayed similar trends to the biological hip joint moment profiles, as expected from the tuning procedure. However, due to individual differences in gait pattern, there were noticeable differences among the generated torques on the inter-participant level. Due to these differences, we chose to present the average torque and power profiles for individual participants rather than averaging across them. The torque profiles generated by the two variants under the constant conditions are shown in Figure 3 for three representative participants. The torque profiles under these conditions for all participants can be found in Supplementary Figures 2–5. Both variants applied extension assistance starting from mid- to late swing, and gradually transitioned to flexion assistance during the stance period. In the NMC-5 profiles, the switch from extension to flexion assistance occurred later in the stance phase. In terms of amplitude, the two variants had similar values in the direction of extension, while NMC-4 generated markedly higher values of flexion torque.

**Figure 3:**
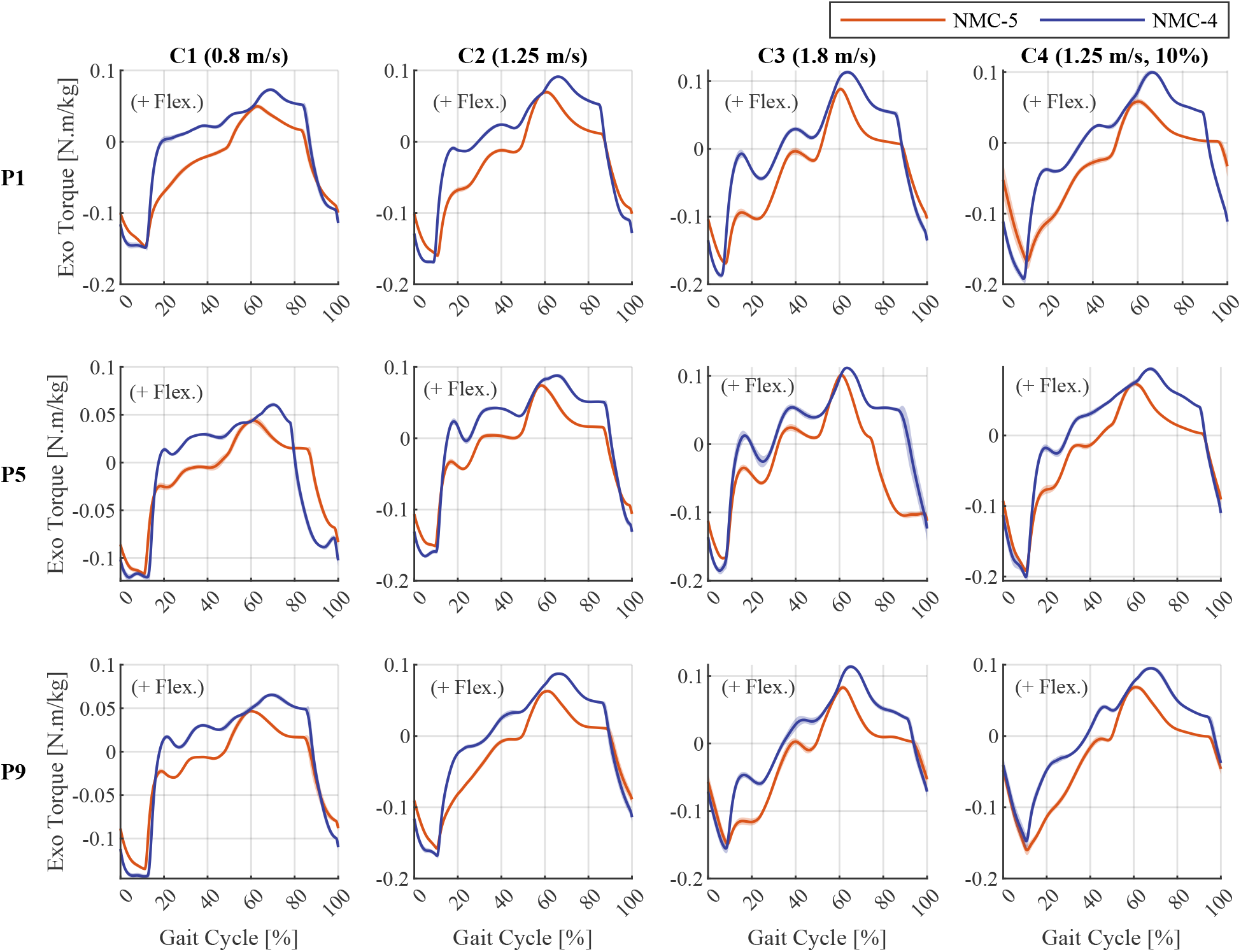
Torque profiles generated by the four-state (NMC-4) and five-state (NMC-5) variants under different conditions (C1–C4) for three representative participants (P1, P5 and P9). The curves represent the mean across strides, and the shaded area represents *±*1 standard deviation around the mean.

The resulting mechanical powers output by the exoskeleton were predominantly positive, as illustrated in Figure 4 for three representative participants in the constant conditions. The power profiles for all participants are provided in Supplementary Figures 6–9 for conditions C1–C4 respectively. In both variants, the most prominent bursts of power occurred around the periods of transition between swing and stance, in accordance with the periods of peak torque observed in Figure 3. However, NMC-4 displayed higher peaks in both positive and negative powers, and also a more oscillatory behavior.

**Figure 4:**
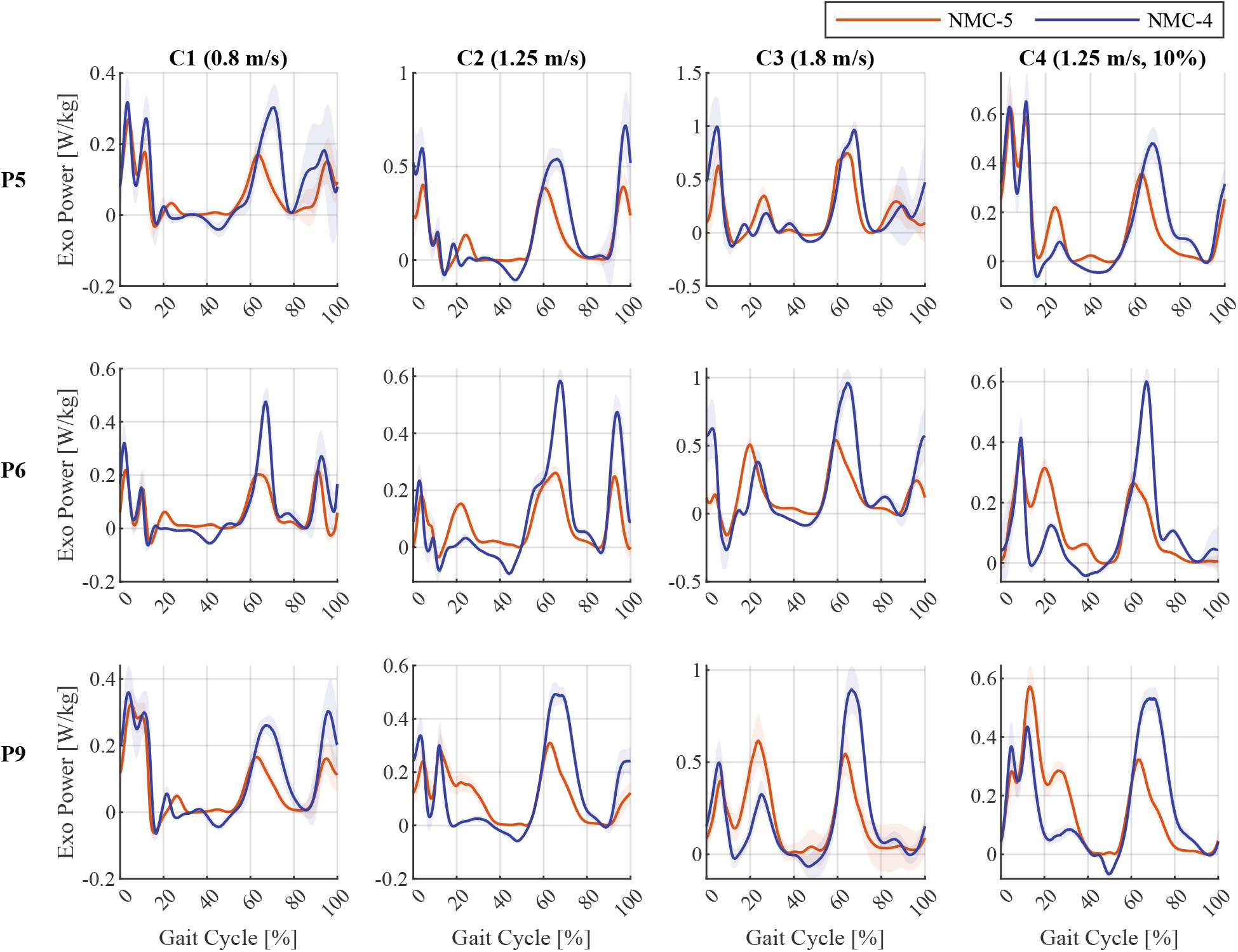
Exoskeleton output mechanical power profiles of the four-state (NMC-4) and five-state (NMC-5) variants under different conditions (C1–C4) for three representative participants (P5, P6 and P9). The curves represent the mean across strides, and the shaded area represents *±*1 standard deviation around the mean.

To better visualize the adaptation of assistance across different walking conditions, the average torque profiles generated by NMC-5 under the constant conditions for each participant are plotted together in Figure 5. While the specific details of the torque profiles varied between participants, the overall trends across the four conditions were largely consistent, particularly in terms of timing.

**Figure 5:**
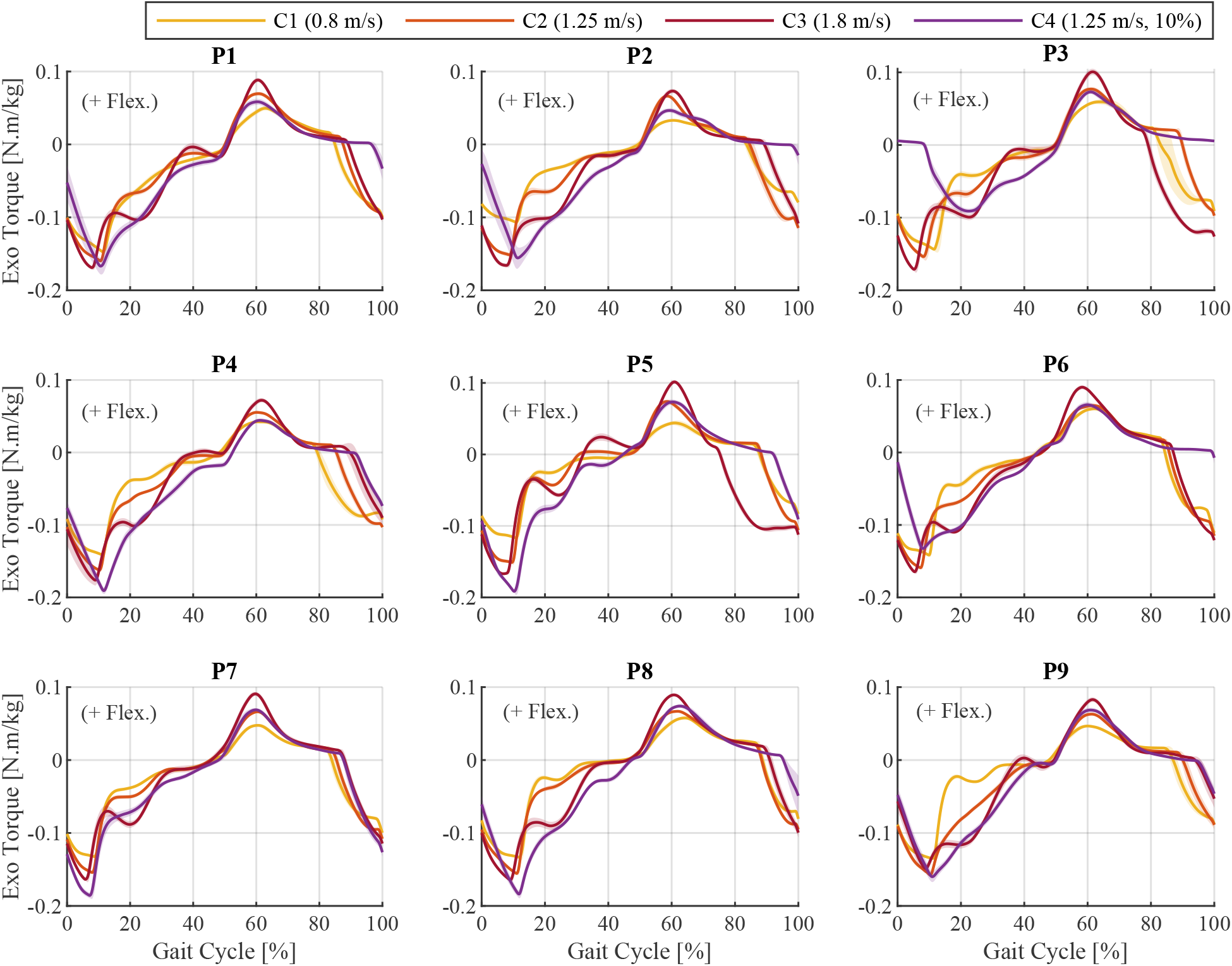
Torque profiles produced by the five-state variant (NMC-5) under the four conditions (C1–C4), shown for each participant (P1–P9) separately. The curves represent the mean across strides, and the shaded area represents *±*1 standard deviation around the mean.

In the variable-inclination condition, the torque profiles displayed a continuous evolution, as illustrated in Figure 6. The evolution was most prominent in the extension assistance period between late swing and mid-stance, where the peak timing increased and the magnitudes decreased with higher inclinations. The unexpected magnitude trends were mostly due to the force-length factor of the virtual muscles, depicted in Supplementary Figure 14.

**Figure 6:**
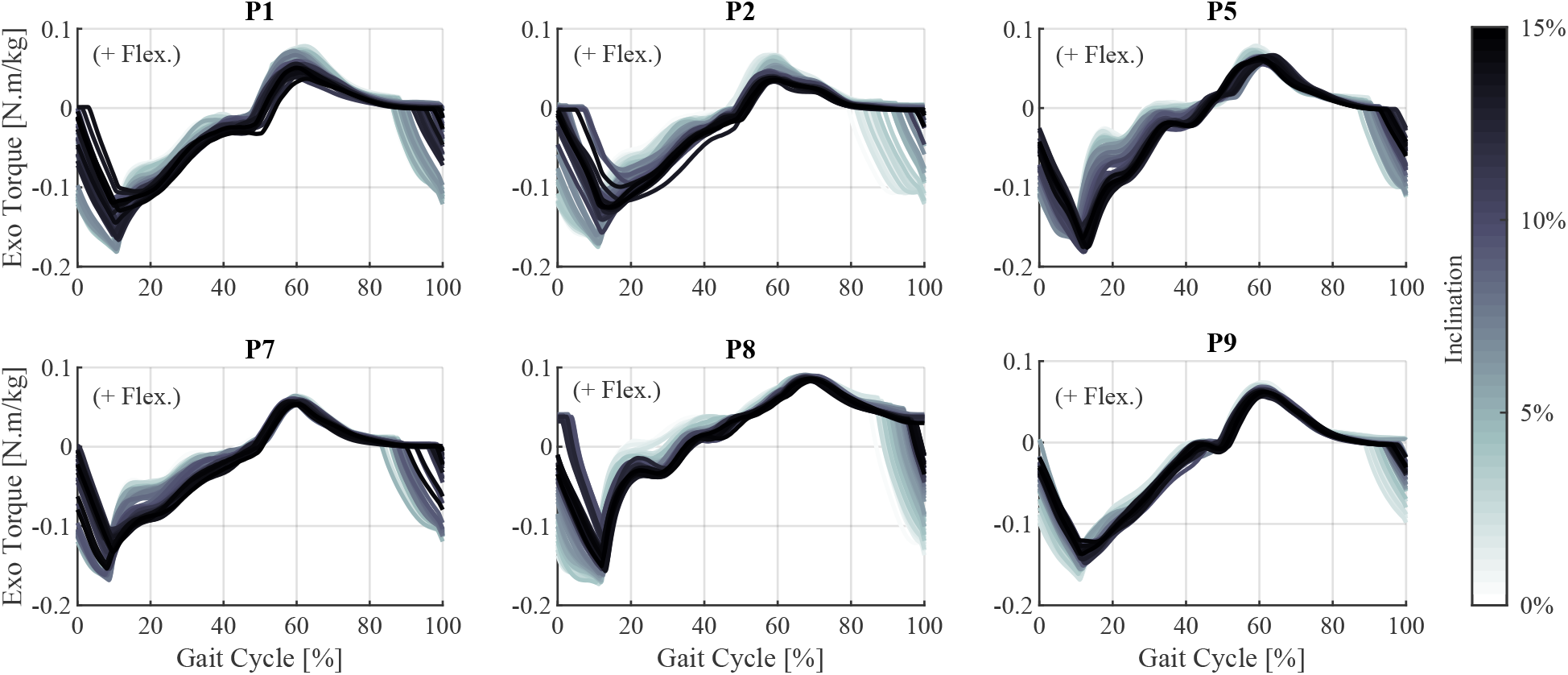
Stride-by-stride evolution of the assistive torque profiles generated by the five-state variant (NMC-5) during the variable-inclination condition (C5) for individual participants. Each curve corresponds to one stride, and the darker curves occurred later in time and therefore correspond to larger inclinations.

### 3.2 Hip joint kinematics

The assistance provided by both variants slightly altered the hip joint angles compared to unassisted walking, as observed in the average profiles in Figure 7A–D. The differences were most prominent in early stance and mid-swing. Furthermore, there were conspicuous differences in the profiles between NMC-4 and NMC-5. The differences were also reflected in the average ranges of motion (RoMs) of the hip joint, plotted in Figure 7E. The hip joint angle profiles for individual participants under different conditions are shown in Supplementary Figures 10–13.

**Figure 7:**
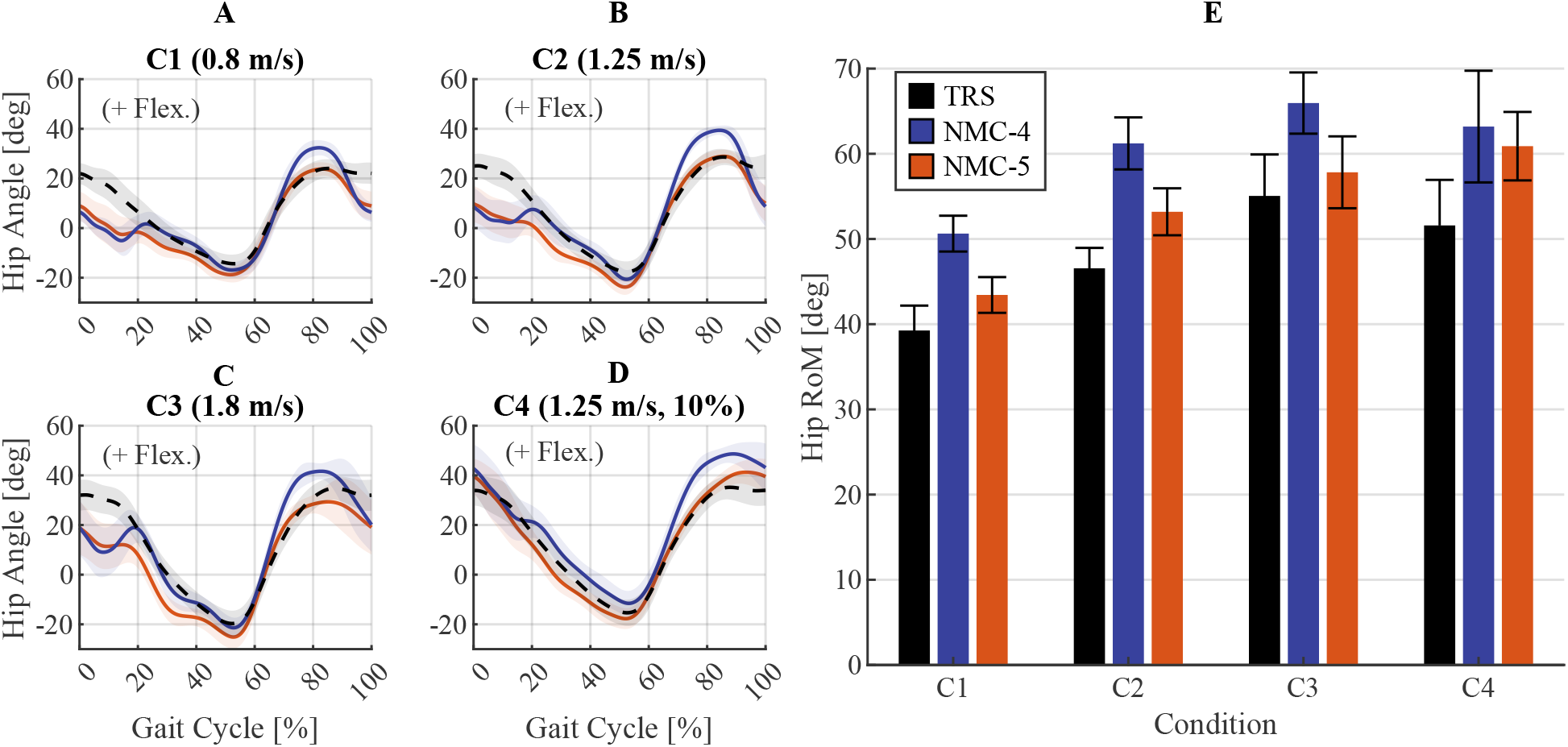
Average hip angle profiles (panels **A**–**D**) and ranges of motion (RoMs) (panel **E**) across all participants obtained with the four-state (NMC-4) and five-state (NMC-5) variants, compared against unassisted walking (TRS). In the average profiles, the curves represent the means across all participants and strides, and the shaded areas mark *±*1 standard deviation around the means.

## 4 Discussion

### 4.1 Generated assistance in the constant conditions

We focus our analysis of the performance of the two variants mostly on the timing of assistance, since the magnitudes were considerably affected by the manually modified gain *G*_*s,v*_. The timings, on the other hand, had a completely emergent nature. When analyzing magnitude, we focus mostly on the relative values within each condition, which were not affected by *G*_*s,v*_.

The FSM state transition timings of both variants were coherent with the gait phase transition timings typically observed in normal walking. Since the transition criteria for the common states between the two variants were identical, the state transition timing values were nearly the same. On average, the states ES, MS-PS, S, and LP started at 0%, 10.26%, 60.6%, and 87.23%, respectively, in accordance with the values reported in normal gait (Winter, 1991).

The generated torque profiles by both variants under each condition were quite similar across different strides for each participant. Contrarily, notable differences in trends and timing were observed among the participants, resulting from individual differences in gait pattern, as observed in Supplementary Figures 10–13. Generally, both variants generated a relatively steep shift toward extension assistance in late swing, leading to a prominent peak of extension torque in early stance. With NMC-4, this peak happened on average at 5.55% in the flat conditions (C1–C3) and at 7.77% in the inclined condition (C4). For NMC-5, the peak occurred around 8.22% in the flat conditions and near 11.02% in the inclined condition. These values are close to the timings observed in both the biological joint moment profiles (Reznick et al., 2021) and those obtained from HitL optimisation (Franks et al., 2022), which occur on average at 6% and 10.3%, respectively. Furthermore, in both variants this peak occurred later as speed and inclination increased, in agreement with the trend observed in biological profiles

The extension-to-flexion torque transition occurring around midstance was remarkably different between the two variants. The transition happened at around 20% to 30% of the gait cycle in case of NMC-4, and had a relatively high variation between the conditions. These values fall in the low range of the biological timings, and precede the ones of HitL optimisation reported at 36.5% *±* 3.5. Furthermore, under the condition C3 (fast walking), the transition from extension to flexion showed an oscillatory behavior for some participants (i.e., P2, P5, P7 and P8 in Supplementary Figure 4), which can disturb the wearer’s movements and lead to discomfort. In contrast, with NMC-5 the transition happened more consistently near 50% of the gait cycle. Thus, NMC-5 provided extension assistance for a longer duration during the stance phase, which can contribute to the forward propulsion of body and reduce the need for ankle push-off.

The peak flexion torques occurred around 67% on average with NMC-4, showing a tendency toward earlier values with increasing speeds. This timing lags behind the typical values of biological and the HitL profiles. With NMC-5, this peak happened around 60% of the gait cycle, and was thus better aligned with the initiation of the swing phase. Moreover, in case of NMC-5, this timing was more consistent across different conditions as evident in Figure 5, in line with the timings in biological torque profiles.

The flexion-to-extension zero crossing happened on average at 88.45% with NMC-4, and tended to occur later with increases in speed or inclination. This is later than the typical timing in biological profiles, but aligns with the ones of the HitL optimisation at 88%. With NMC-5, this timing was around 85.67% of the gait cycle, closer to the average value of 80% observed in biological profiles. For both variants, this timing showed the highest inter-participant variability. This is due to the dependence of this timing on the initiation of the LP state. The switch from S to LP is the only state transition driven by the angular velocity signal (rather than ground contact, which is used for the rest of the state transitions), and is thus more sensitive to individual gait patterns.

Overall, in terms of timing, NMC-5 showed better alignment with both the biological and the HitL profiles. Notably, in case of NMC-4, the earlier zero crossing from extension to flexion and the later flexion to extension transition resulted in a rather short extension assistance duration, lasting around 40% of the gait cycle. This duration was closer to 60% with NMC-5, showing higher coherence with the duration of the stance phase. Additionally, the direct dependence of the length feedback used in the MS-PS state of NMC-4 on hip angle resulted in oscillation of the torques between positive and negative values in mid-stance for some participants. This aspect was improved in NMC-5 thanks to the separation of the MS and the PS states, which allowed more control over tuning of the reflexes.

In terms of magnitude, the average peak extension to flexion ratio in NMC-4 profiles was 1.71, with a decreasing trend at higher speeds and inclinations; the maximum ratio of 1.91 was obtained in the C1 (slow walking) condition, and the minimum value of 1.66 was observed in C4 (medium speed at 10% inclination). This value is 40% lower than the ratio observed for NMC-5, namely 2.45 on average. In biological profiles, this ratio is above 1.5, and around 2.3 in the HitL profiles, and has an increasing trend with faster speeds and higher inclinations. A similar increasing tendency was observed in the NMC-5 profiles for most participants at higher speeds and in the inclined condition. Furthermore, the absolute extension peak in NMC-4 profiles increased with speed but decreased with the ground inclination, although a higher effort is required in uphill walking. In contrast, NMC-5 generated higher extension peaks in C4 compared to C2 for all participants except two (P3 and P6, as shown in Figure 5).

Concerning mechanical powers and works, both variants resulted in mostly positive work (0.158 versus *−*0.011 J *·* kg^*−*1^ for NMC-4, and 0.114 versus *−*0.002 J *·* kg^*−*1^ for NMC-5). This indicates that the applied torques were mostly aligned with the user’s intended movement, since the torques were not high enough to enforce the movements. NMC-4 applied higher peak powers in both positive and negative directions as seen in Figure 4, resulting in 1.43 times more positive and 13.36 times more negative mechanical work per stride than NMC-5 on average. A major proportion of the positive power delivered by NMC-4 occurred during swing, resulting in a visible overshoot of the swing leg, as can be observed in Figure 7A–D. This added positive power was therefore not efficient in contributing to the forward propulsion of the user. The majority of the negative work performed by NMC-4 was produced in the MS-PS state, because of the early transition to flexion torques as discussed earlier. For NMC-5, the minor negative work was mostly due to fluctuations in the hip angular velocity measured by the actuators of the exoskeleton around 10–20% of the gait cycle, which were artefacts of the movement of the attachments relative to the body, as will be discussed in Section 4.2. For both variants, the negative work increased at higher walking speeds, and was the lowest in the inclined condition.

### 4.2 Effects on hip joint kinematics

Both variants extended the hip range of motion in all conditions compared to the transparent trial. NMC-4 increased the range of motion in flexion mostly, by an average of 9.90^*°*^ in C1, 13.26^*°*^ in C2, 11.19^*°*^ in C3, and 7.80^*°*^ in C4. Inversely, NMC-5 caused higher extension angles with increases of 3.50^*°*^ in C1, 5.18^*°*^ in C2, 5.59^*°*^ in C3, and 4.97^*°*^ in C4 compared to unassisted walking. These results are in line with the patterns observed in the torque and power profiles of the two variants; namely, the higher magnitudes and longer durations of flexion assistance by NMC-4, and the sustained extension assistance during mid- and late stance by NMC-5. Note that although NMC-4 led to a bigger increase in the range of motion, this increase was in large part due to overshooting of the swing leg. This is evident by comparing the final hip angle at the moment of heel-strike, which was similar between the two variants. Therefore, this added movement did not change the effective range of motion since it did not lead to a longer step. In contrast, the increase induced by NMC-5 resulted from extending the stance leg further, thus pushing the body further forward.

The hip angle profiles exhibited an abnormal oscillatory pattern in early to mid-stance, particularly in the flat walking conditions (C1–C3). This pattern was more pronounced with NMC-4, as evident in Figure 7A–C. By inspecting video recordings of the trials, this pattern was found to be largely an artefact of the relative movement of the exoskeleton interfaces with respect to the user’s body. These movements happened due to compliance in the interface materials and the imperfect fitting of the attachments which allow a slight play at the interface. Therefore, this phenomenon coincided with the peak of extension torque, and was aggravated by the subsequent steep transition to flexion torque with NMC-4. Besides the measurement artefact, this behavior can also lead to discomfort and negatively affect the forward momentum of the user.

### 4.3 Adaptation of the assistance to varying inclination

The NMC-5 torques showed a gradual and continuous adaptation to the changes in the inclination, as evidenced by Figure 5. In terms of trend, the adaptation had a similar characteristic pattern across participants. The adaptation trend in terms of timing had similarities to the trends observed in biological torque profiles at different inclination levels (Winter, 1991; Perry et al., 1992; Reznick et al., 2021). Namely, the extension assistance period shifted to the right (i.e., later in the gait cycle) as the inclination increased. As a result, the duration of extension assistance during the stance phase increased for higher slopes, shifting gradually from an average of 44.89% of the gait cycle for flat walking to 48.9% at 15% inclination. Also, the extension peak in stance happened later as the inclination increased, shifting gradually from an average of 8.41% during flat walking to 12.02% for the 15% inclination.The flexion peak during swing did not undergo major shifting in time, even though the duration of flexion assistance increased at higher inclinations, and the transition to extension torque tended to happen later. These results are in accordance with the previously referenced biological torque profiles.

On the other hand, the trends observed in torque magnitudes did not completely follow the biological torque adaptation patterns. In particular, the extension assistance magnitude tended to slightly decrease at higher slopes, with a reduction of around 20% at the final inclination (15%) compared to level walking. This is against the adaptation observed in biological torques, as walking on steeper inclinations requires more effort. The flexion peak also had a decreasing tendency, reducing by 24% at the final inclination compared to flat walking. However, the reducing pattern of flexion peaks is coherent with biological profile trends (Winter, 1991; Perry et al., 1992; Reznick et al., 2021). Both of these reductions at higher inclinations are related to the more flexed posture at the hip in higher inclinations. The reduction in extension and the increase in flexion angles lead to a lower CE length for GLU, and inversely to a higher CE length range for ILPS. The effect of these changes in CE length is lower values of the force-length factor, *f*_*l*_ (Equation 6), since the optimal length (*l*_*opt*_) is tuned based on the CE length range in level-ground walking conditions. As a result, at higher inclination levels, the output of the force-length relationship decreases near the beginning and end of the gait cycle, as shown in Supplementary Figure 14. The generated torque values are therefore reduced, particularly in case of the extension peak which happens in early stance. It is worth reminding that in condition C5, the manually tuned gain *G*_*s,v*_ was kept constant, and therefore the observed changes in magnitude were entirely driven by the changes in controller inputs.

### 4.4 Limitations and future directions

In this work, we focused on the technical validation and assessment of the novel NMC structure. Therefore, the main outcomes of interest were the outputs of the controller. Future studies evaluating more direct functional outcomes of the assistance, such as reductions in metabolic energy expenditure or muscular activities of users, are needed for a more comprehensive assessment of the practical benefits. For such studies, parameter tuning should also be purely guided by the targeted functional outcomes, rather than mimicry of the biological torque profiles, which was used as one of our major metrics for this study. However, given the relatively large number of parameters, the tuning procedure remains difficult. Even though the approach proposed in this study based on the roles of major parameters and separate stages can speed up the procedure, a more systematic approach would be highly beneficial.

In terms of functionality, the main shortcoming in both of the tested variants was the insufficiency of magnitude adaptation of the torques to changes in walking speed and ground inclination. An intuitive way to address this issue is using variable reflex gains as a function of speed and inclination. However, such a solution would come at the cost of requiring speed and inclination estimators. Another option would be to design the force-length and force-velocity functions of the virtual muscles based on the desired behavior, rather than using the relationships used in biological muscle modeling. This approach can also be used to make the virtual muscles complement their biological counterparts, by delivering higher forces in configurations where the biological muscles have reduced capacity or efficiency.

## 5 Conclusion

This study presented a novel structure for NMCs, with modifications to the virtual muscle model and reflex modulation. Based on the proposed general structure, two controller variants for hip exoskeletons were proposed, with four- and five-state modulation of reflexes (NMC-4 and NMC-5). For setting the controller parameters, a sequential and iterative approach along with data-driven simulation methods and leveraging key parameter identification allowed a targeted tuning without the need for costly optimization procedures. In validation experiments, both variants demonstrated adaptability of the assistance torque profile as a function of the gait pattern. The adaptations led to differences in the characteristic timings, flexion and extension peak magnitudes and magnitude ratios across participants and conditions. A higher inter-participant variability (compared to the stride-to-stride variability for each individual) corroborated the potential of personalized assistance using NMCs. The adaptation of the assistance to different walking conditions in terms of timing emerged directly from the interplay between the neuromuscular model and the users’ gait pattern adaptations. The adaptation of magnitudes, on the other hand, had to be improved using a condition-dependent scaling gain that was adjusted manually. The five-state variant showed a more desirable adaptive behavior both in terms of timing and relative magnitudes, underlining the utility of a more fine-grained reflex modulation.

## Acknowledgments

We would like to thank Dr Romain Baud and Dr Olivier Pajot for their contributions to the preparation of the software and hardware of the exoskeleton, and Olivier Clerc for his assistance in carrying out the experiments.

## Data Availability Statement

The data collected in the experiments are available from the corresponding author upon reasonable request.

## Ethical Standards

The authors assert that all procedures contributing to this work comply with the ethical standards of the relevant national and institutional committees on human experimentation and with the Helsinki Declaration of 1975, as revised in 2008.

## Author Contributions

Methodology and Investigation: A.R.M; S.M; A.D.R. Software: S.M; Formal Analysis and Visualization: A.R.M; S.M. Supervision and Funding Acquisition: A.I; M.B. Writing – Original Draft: A.R.M; S.M. Writing – Review & Editing: A.I; M.B. All authors participated in the conceptualization of the study and approved the final submitted draft.

## Footnotes

3 The subscript “*nlp*” is replaced either by “*se*”, “*pe*” or “*be*” in the equations for each element.

4 The equivalent notations in (Geyer and Herr, 2010) are *l*_*se,slack*_ = *l*_*rest*_, *l*_*be,slack*_ = *l*_*min*_, *l*_*pe,slack*_ = *l*_*opt*_.

5 The equivalent notations in (Geyer and Herr, 2010) are 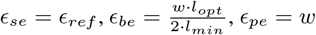.

