## Supplementary material for "Novel Neuromuscular Controllers with Simplified Muscle Model and Enhanced Reflex Modulation: A Comparative Study in Hip Exoskeletons": Supplemetary Material

PREPRINT VERSION

**Ali Reza Manzoori**[*]
Biorobotics Laboratory
EPFL
Lausanne, CH-1015

**Sara Messara** [†]
Biorobotics Laboratory
EPFL
Lausanne, CH-1015

**Andrea Di Russo**
Biorobotics Laboratory
EPFL
Lausanne, CH-1015

**Auke Ijspeert**
Biorobotics Laboratory
EPFL
Lausanne, CH-1015

**Mohamed Bouri**
Biorobotics Laboratory & Translational Neural Engineering Laboratory
EPFL
Lausanne, CH-1015

May 10, 2024

| Parameter [Unit] | GLU value | ILPS value |
|---|---|---|
| $l_{slack}[m]$ | 0.1 | 0.1 |
| $l_{opt}[m]$ | 0.157 | 0.13 |
| $\phi_{ref}[deg]$ | 170.0 | 180.0 |
| $F_{max,ce}[N]$ | 1500 | 1500 |
| $v_{max}[N]$ | 1.32 | 1.32 |
| $\tau[s]$ | 0.1 | 0.01 |
| $w[]$ | 0.56 | 0.56 |
| $\phi_{max}[deg]$ | 0 | 0 |
| $\phi_{off}[deg]$ | 0 | 0 |
| $r_0[m]$ | 0.1 | 0.1 |
| $\rho[]$ | 0.5 | 0.5 |
| $l_{off}[m]$ | 0.157 | 0.157 |
| $K[]$ | 0.005 | 0.005 |
| $N[]$ | 1.5 | 1.5 |
| $C[]$ | 0.05 | 0.05 |
| $\delta t[s]$ | 0.005 | 0.005 |
| $Stim_0[]$ | 0.01 | 0.01 |
| $\delta_{stim}[]$ | 0.25 | 0.25 |

Supplementary Table 1: Muscle model parameter values.

[*]Co-first author
[†]Co-first author

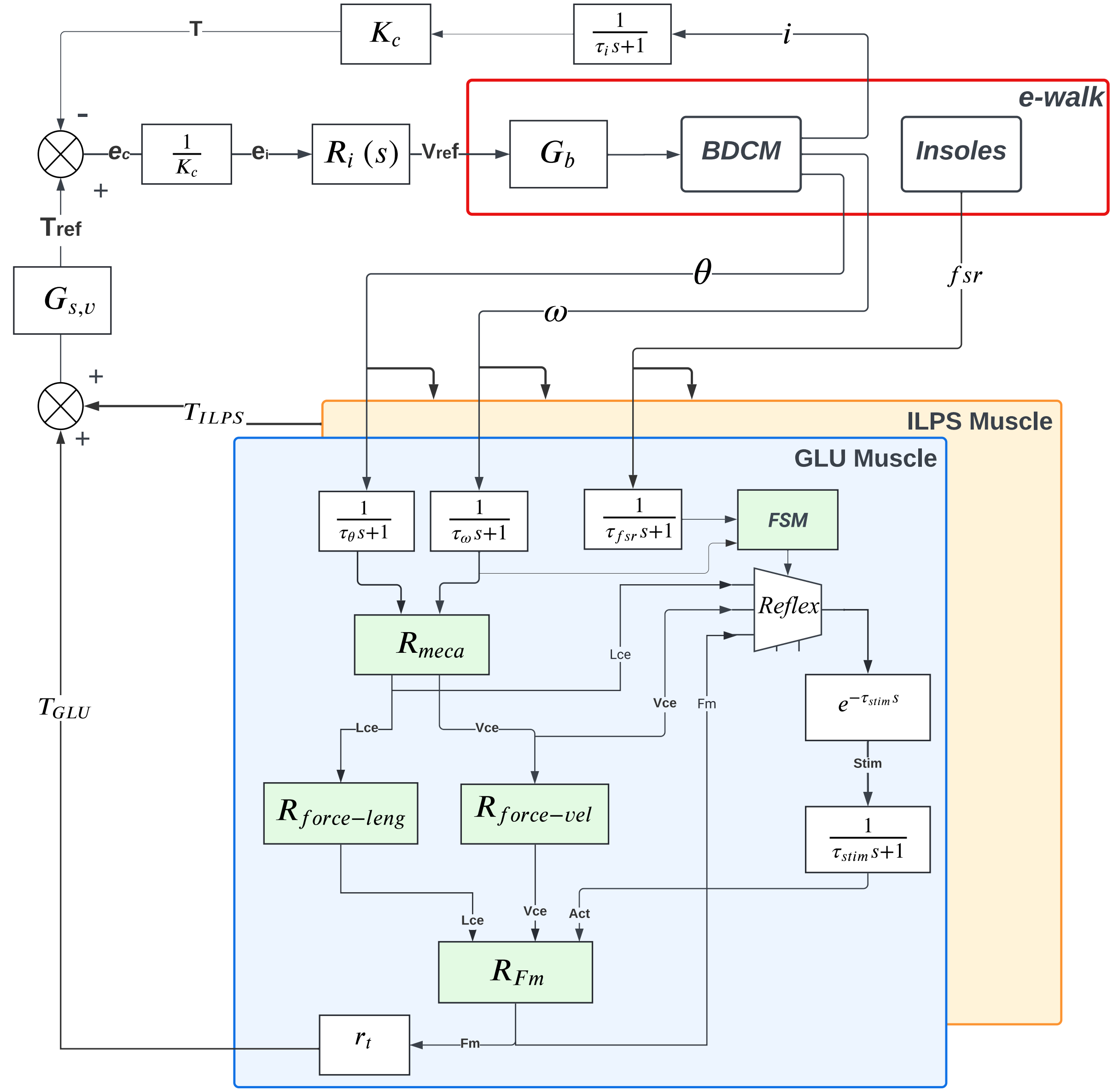


Supplementary Figure 1: Detailed block diagram of the novel NMC architecture, as implemented on the eWalk hip exoskeleton.

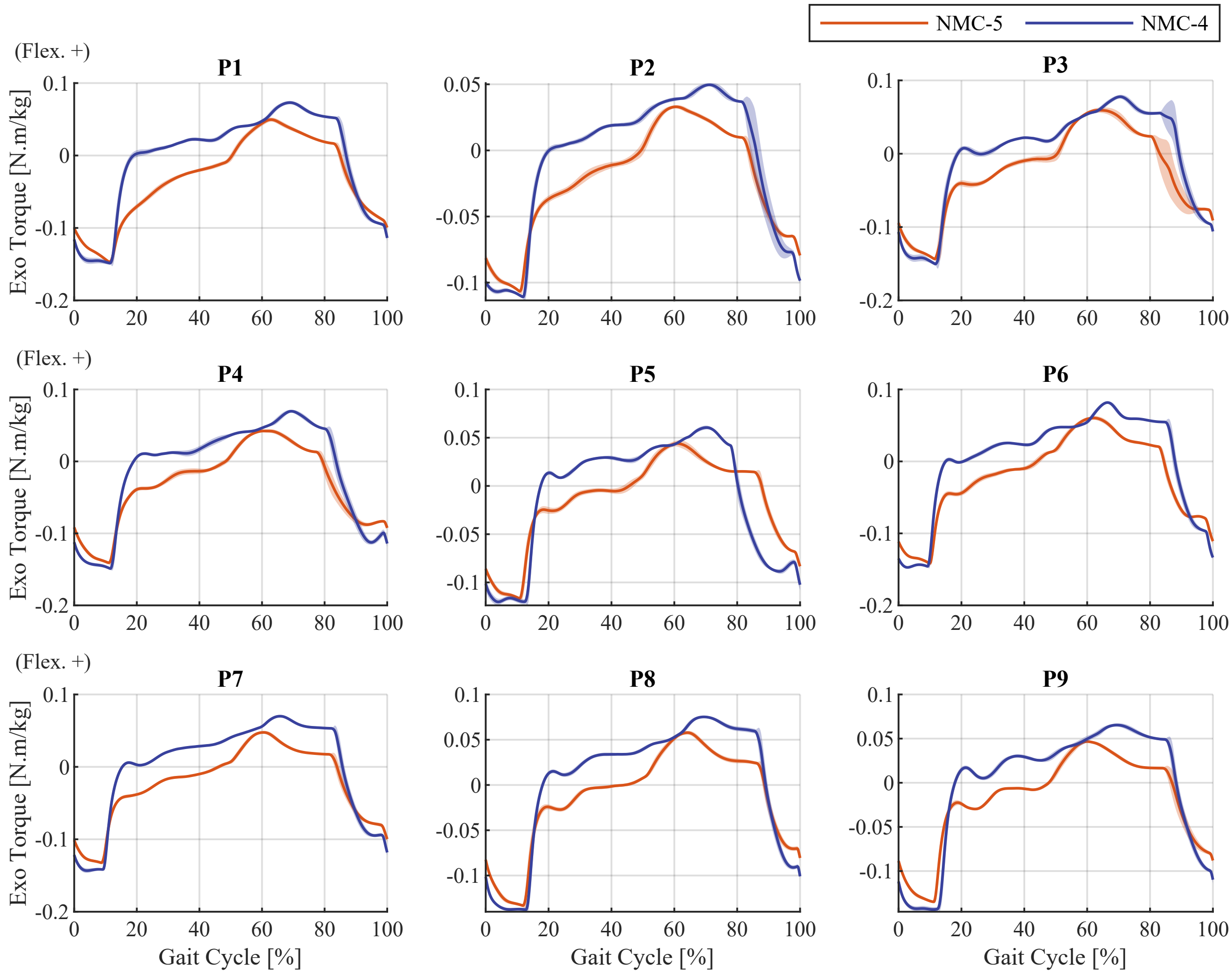


Supplementary Figure 2: Torque profiles generated by the four-state (NMC-4) and five-state (NMC-5) variants under condition C1 for all participants.

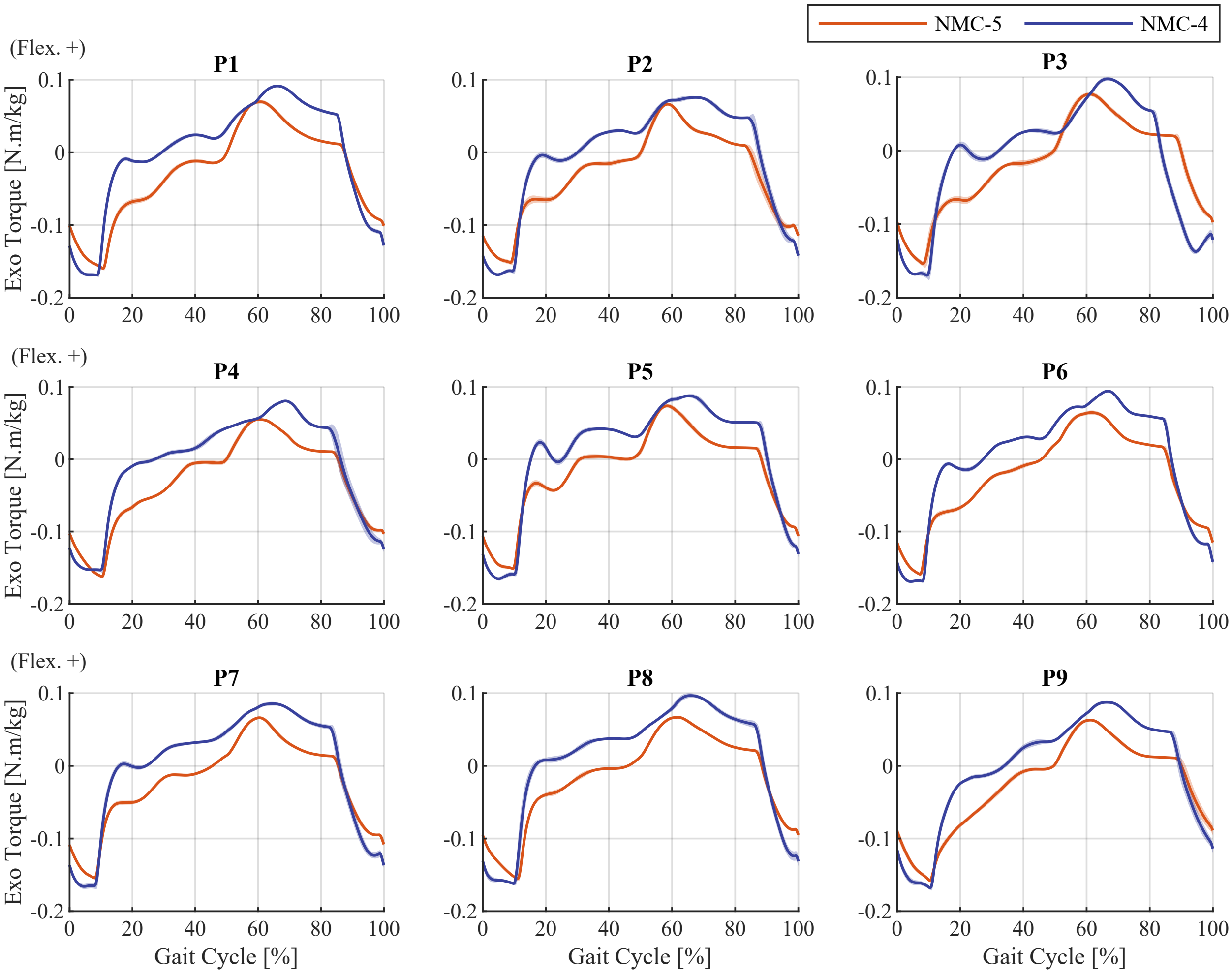


Supplementary Figure 3: Torque profiles generated by the four-state (NMC-4) and five-state (NMC-5) variants under condition C2 for all participants.

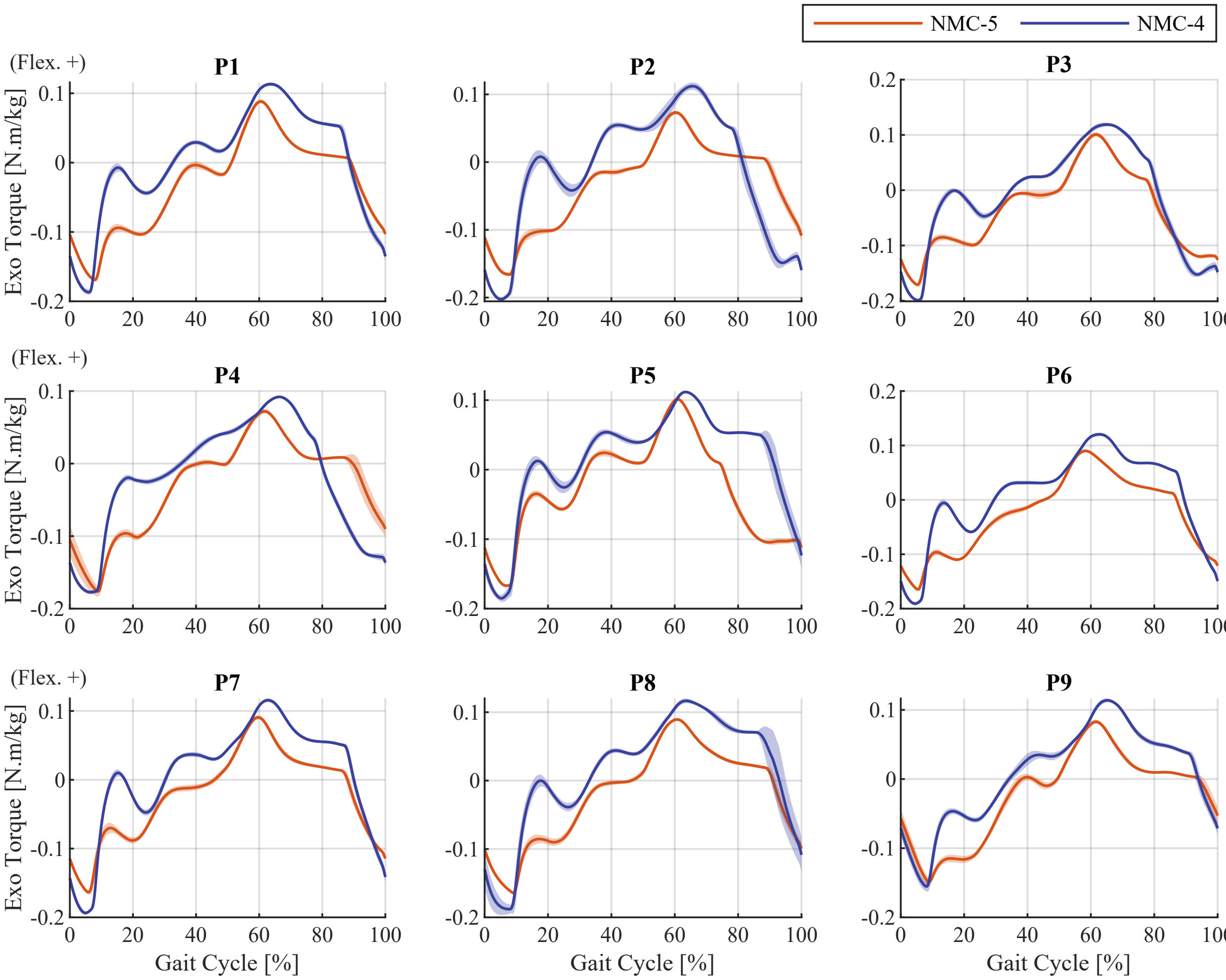


Supplementary Figure 4: Torque profiles generated by the four-state (NMC-4) and five-state (NMC-5) variants under condition C3 for all participants.

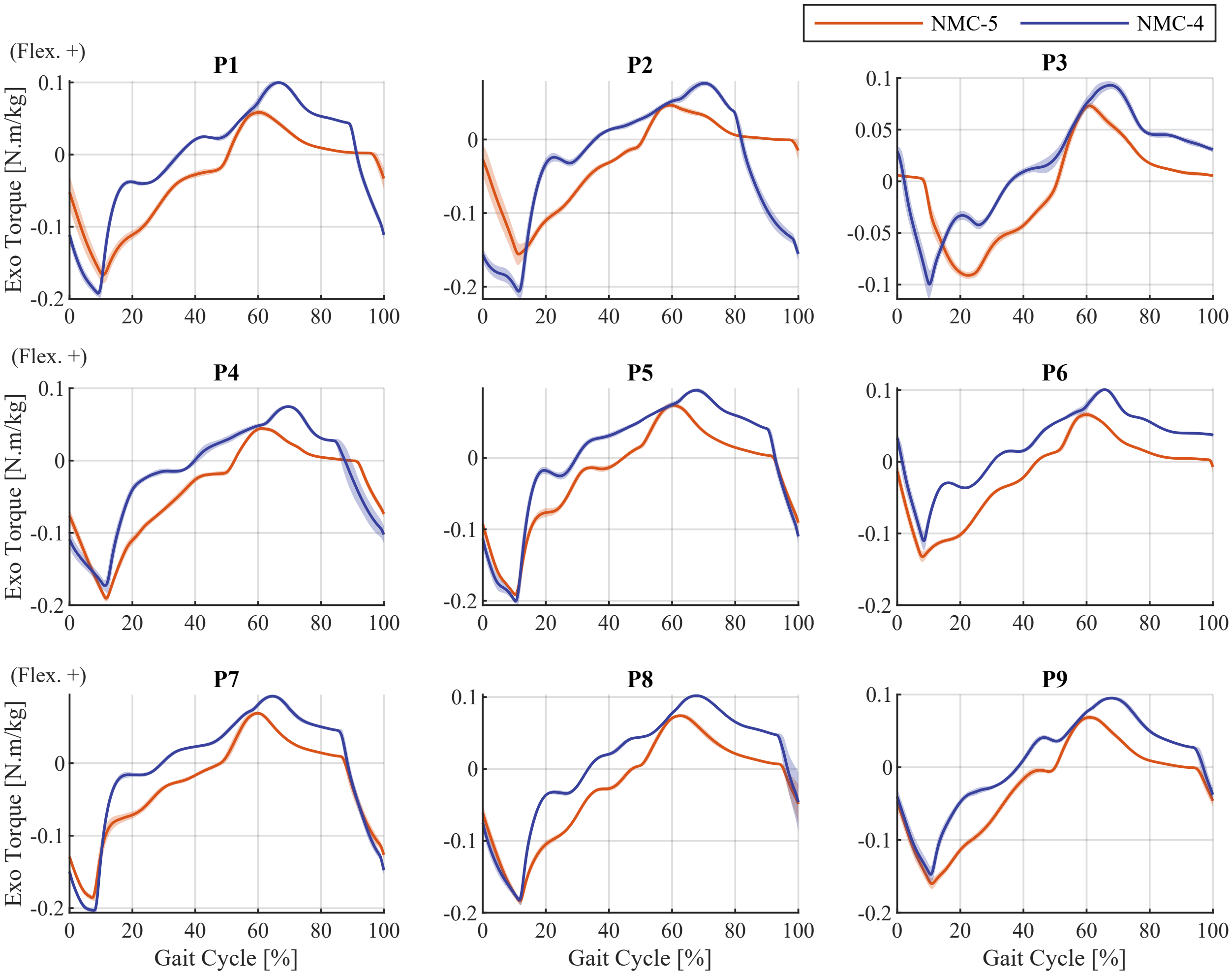


Supplementary Figure 5: Torque profiles generated by the four-state (NMC-4) and five-state (NMC-5) variants under condition C4 for all participants.

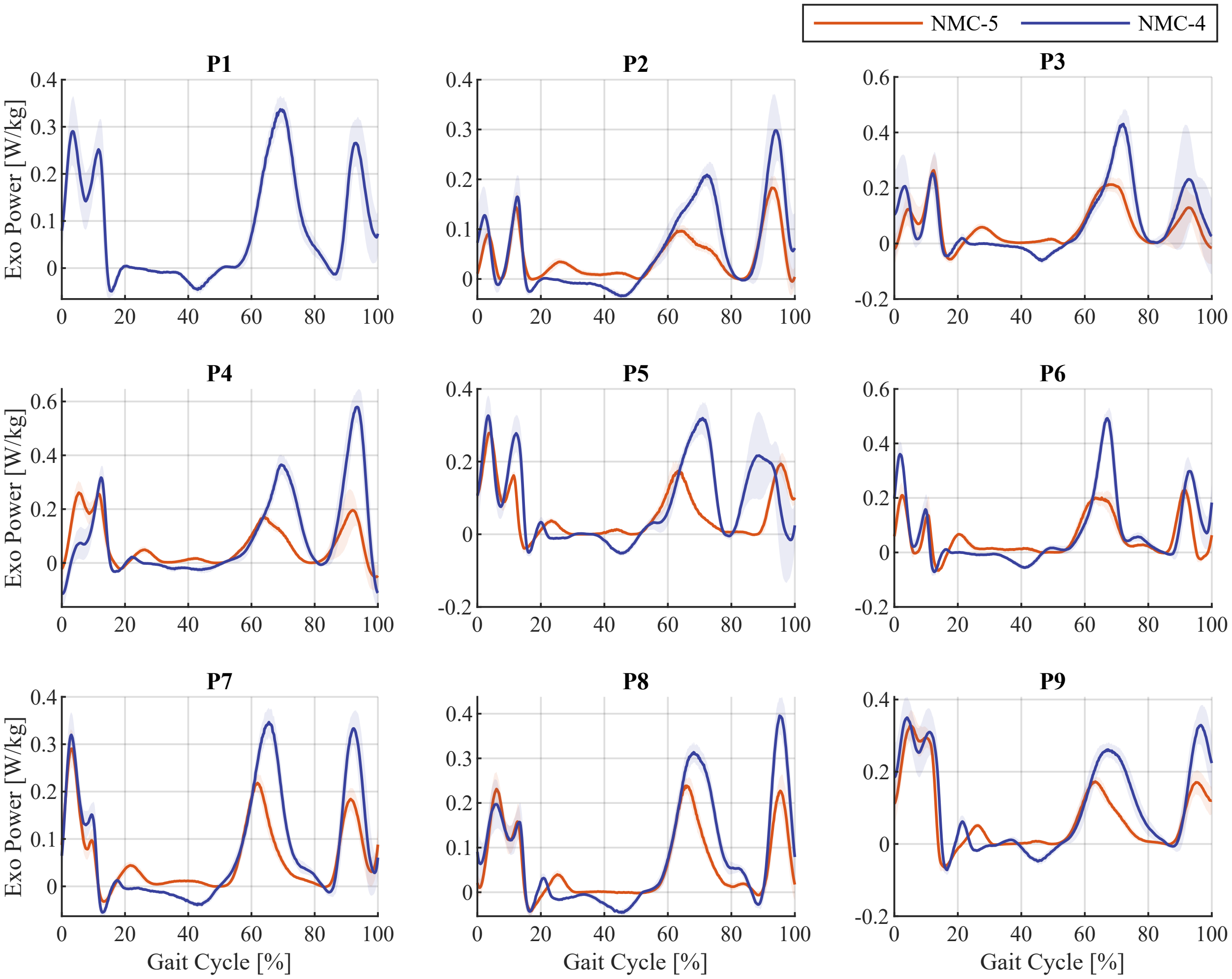


Supplementary Figure 6: Exoskeleton mechanical power output profiles with the four-state (NMC-4) and five-state (NMC-5) variants under condition C1 for all participants. Note that for participant P1, due to an issue in the logging system, the power profile for NMC-5 was not available.

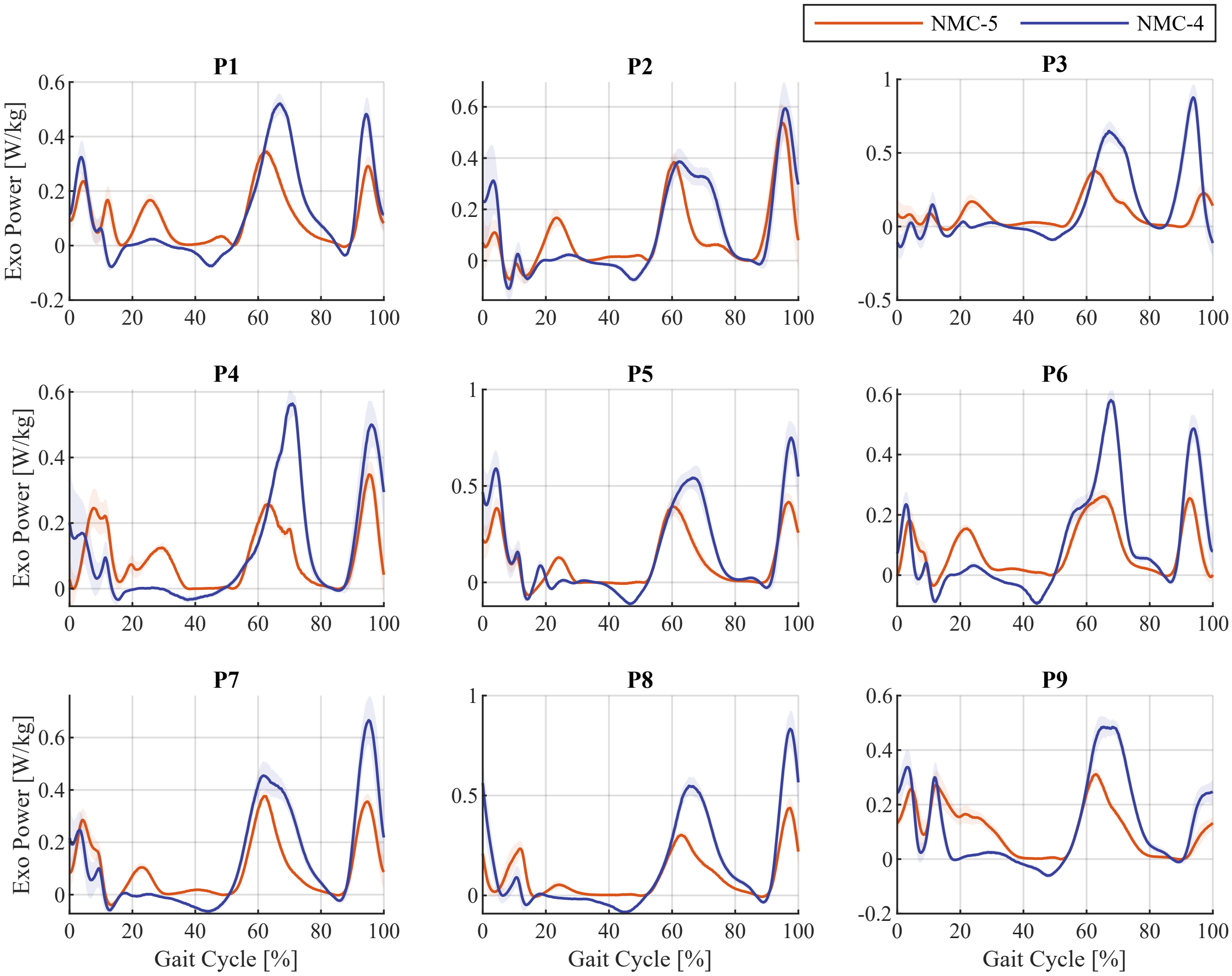


Supplementary Figure 7: Exoskeleton mechanical power output profiles with the four-state (NMC-4) and five-state (NMC-5) variants under condition C2 for all participants.

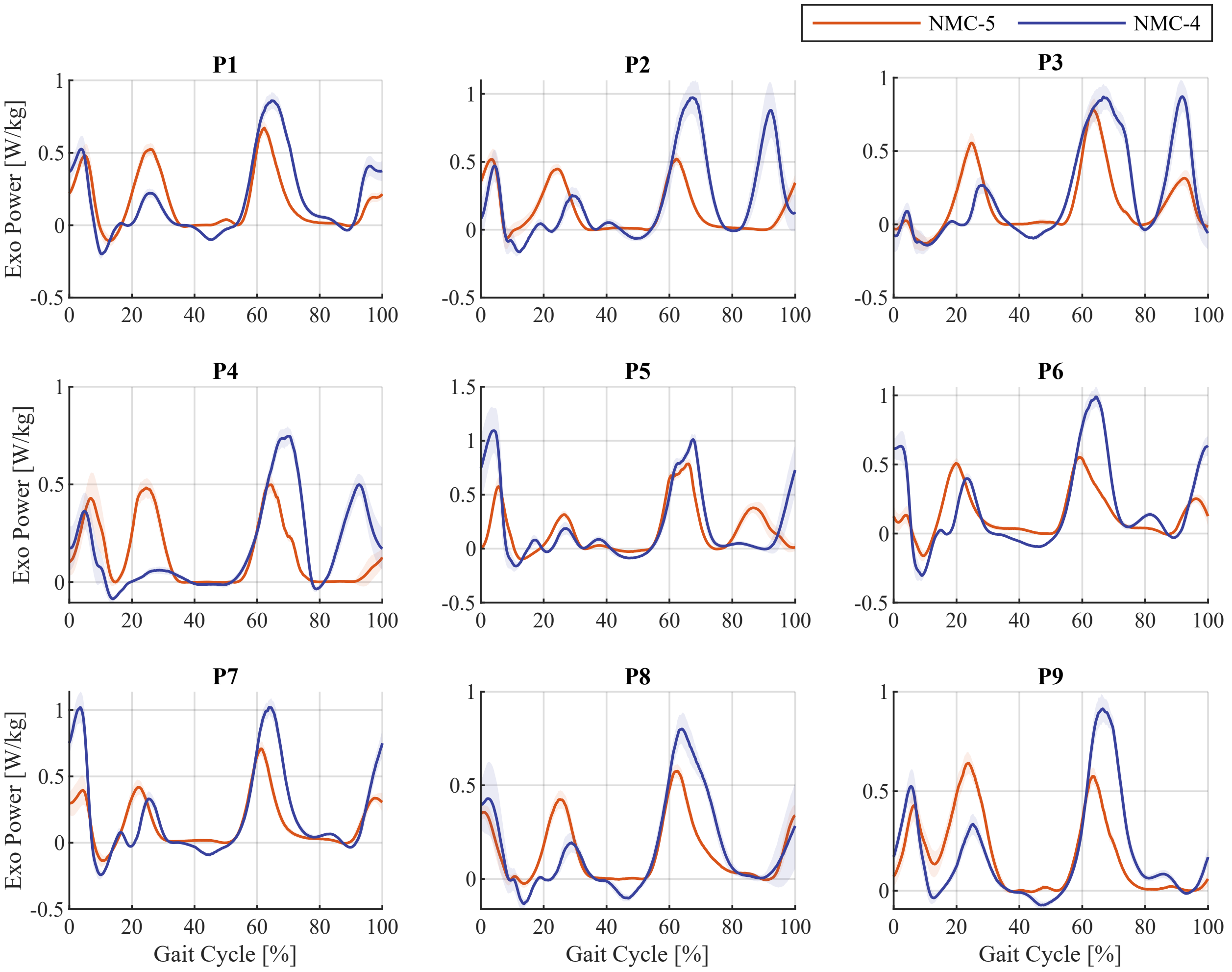


Supplementary Figure 8: Exoskeleton mechanical power output profiles with the four-state (NMC-4) and five-state (NMC-5) variants under condition C3 for all participants.

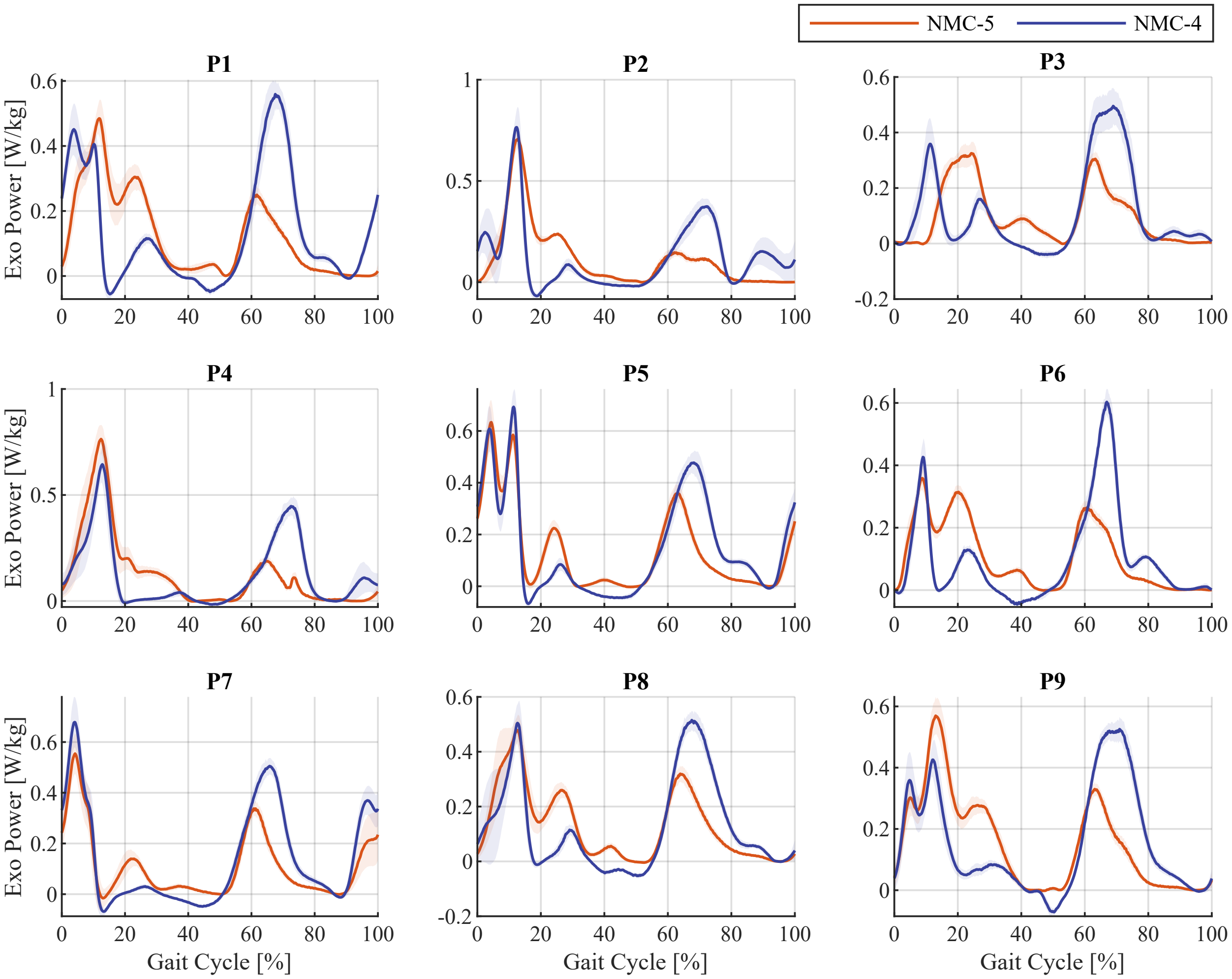


Supplementary Figure 9: Average exoskeleton mechanical power output profiles with the four-state (NMC-4) and five-state (NMC-5) variants under condition C4 for all participants.

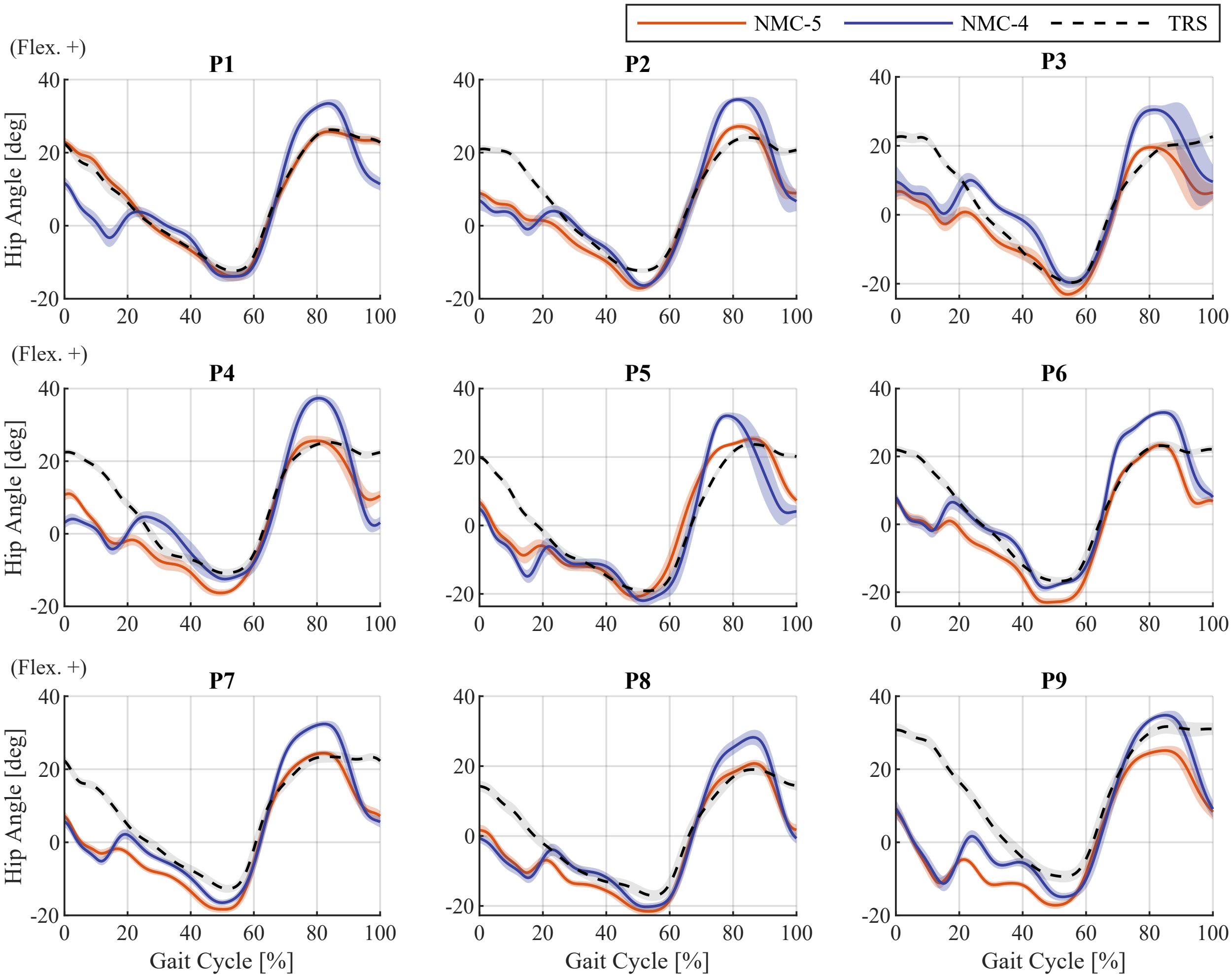


Supplementary Figure 10: Average hip angle profiles in assisted walking (NMC-4 and NMC-5) and unassisted mode (TRS) for all participants under condition C1.

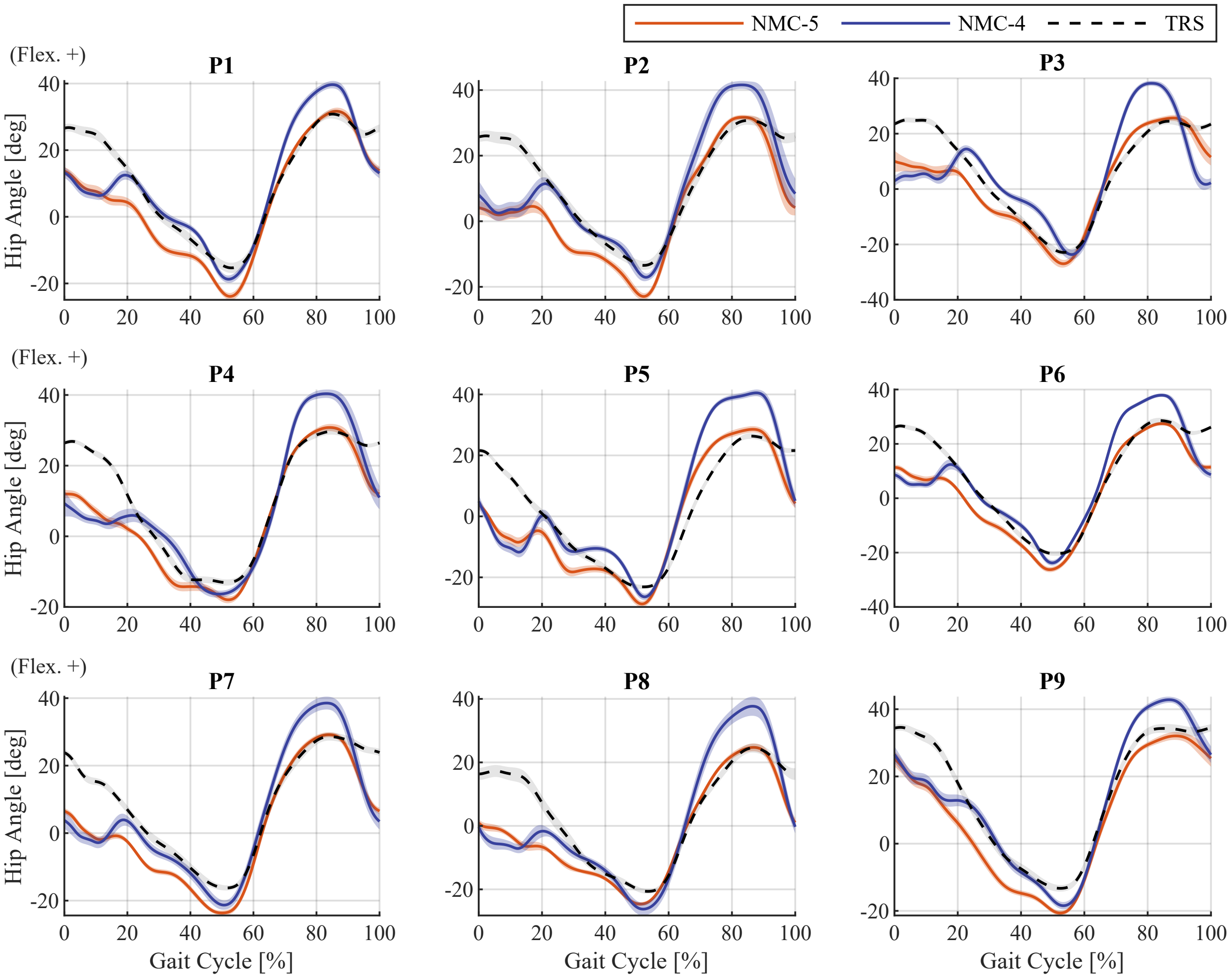


Supplementary Figure 11: Average hip angle profiles in assisted walking (NMC-4 and NMC-5) and unassisted mode (TRS) for all participants under condition C2.

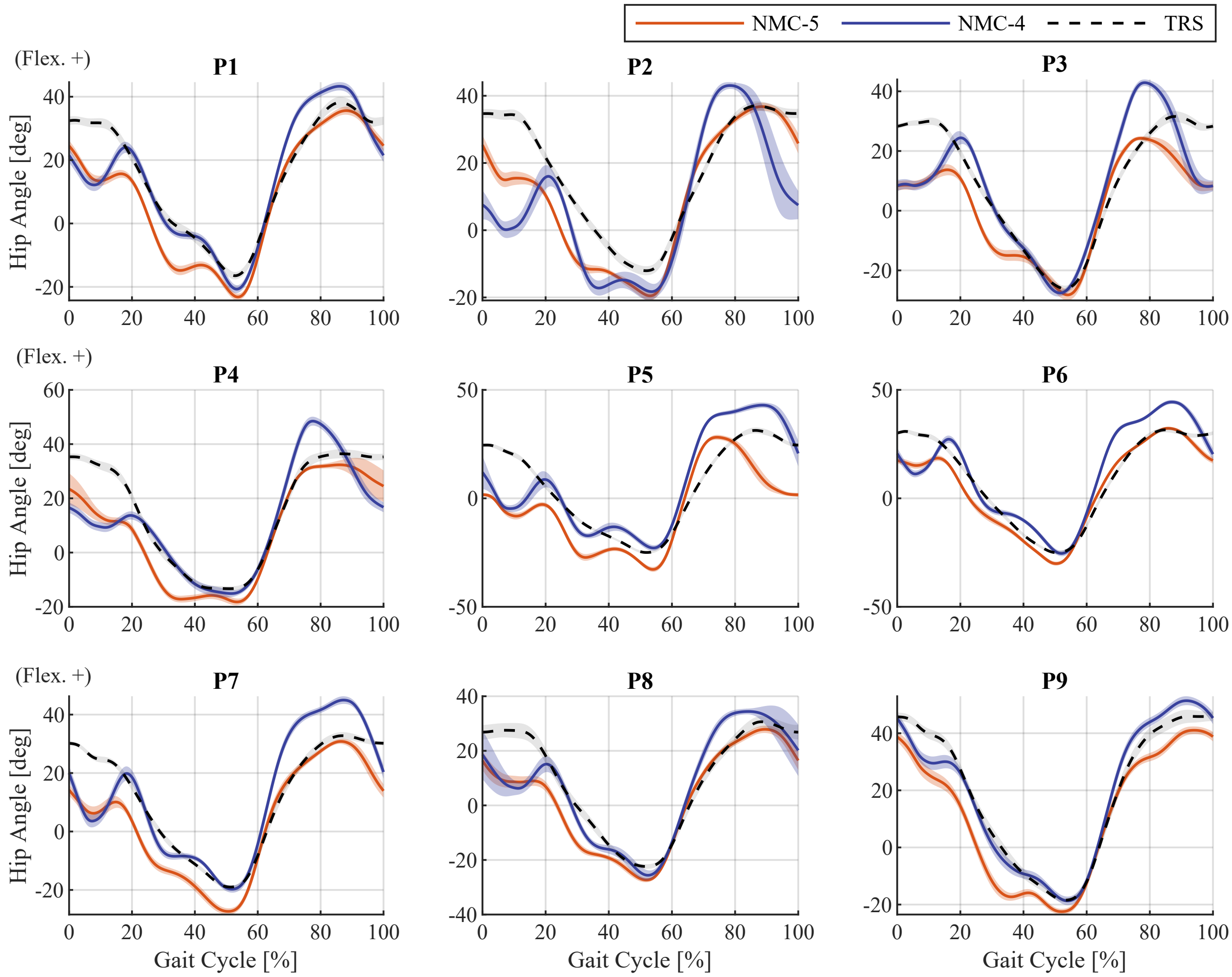


Supplementary Figure 12: Average hip angle profiles in assisted walking (NMC-4 and NMC-5) and unassisted mode (TRS) for all participants under condition C3.

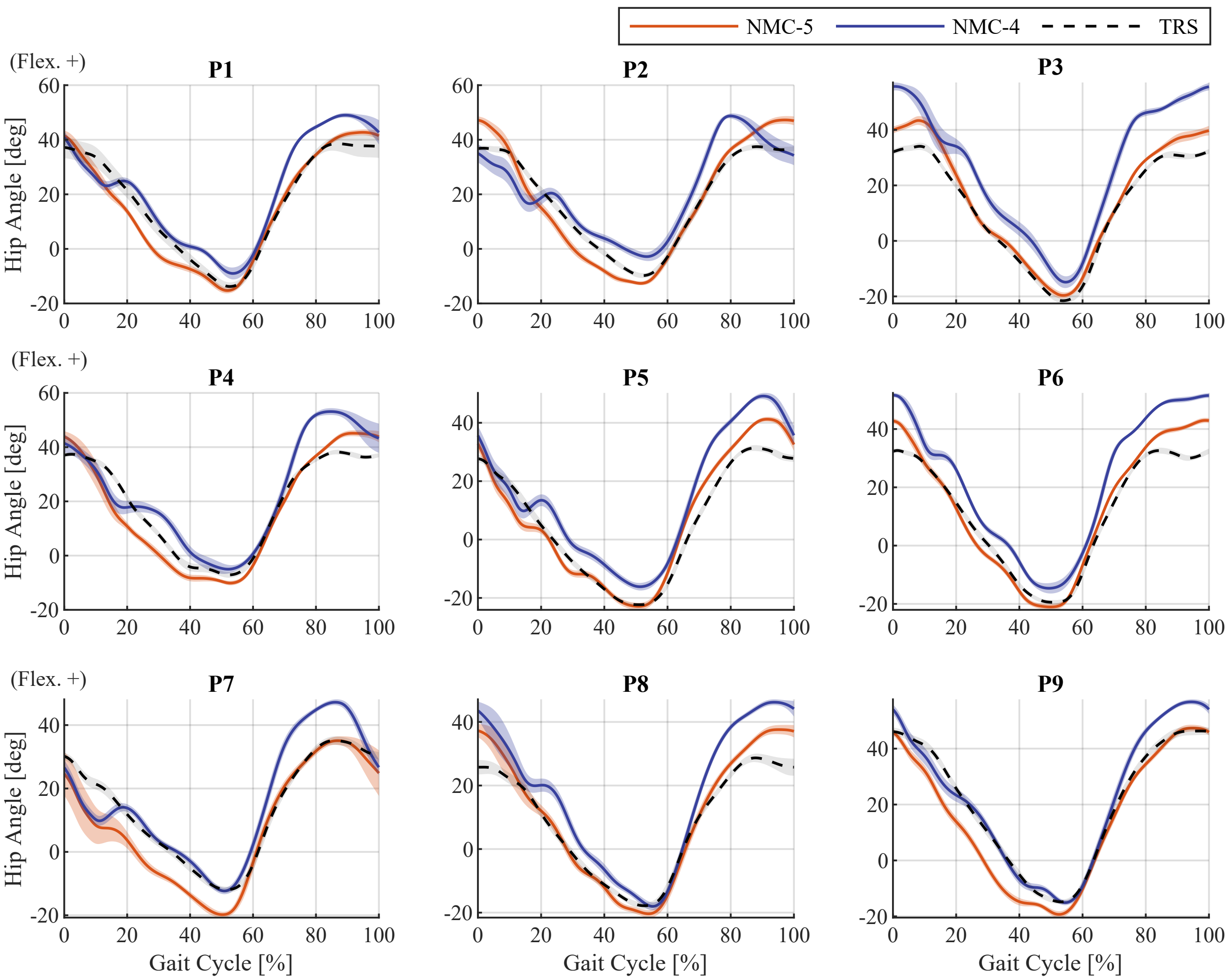


Supplementary Figure 13: Average hip angle profiles in assisted walking (NMC-4 and NMC-5) and unassisted mode (TRS) for all participants under condition C4.

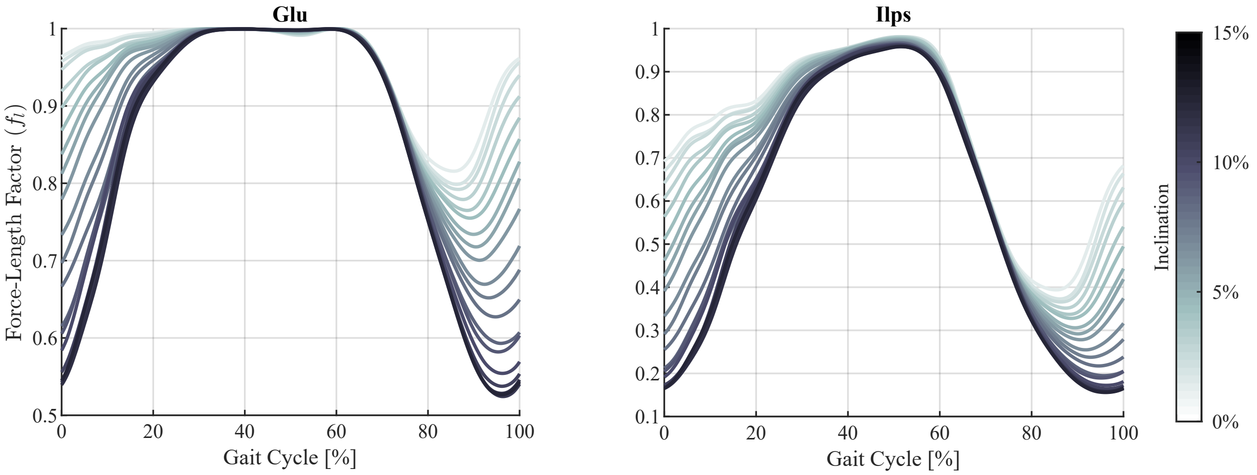


Supplementary Figure 14: Evolution of the force-length factor ($f_l$) for the extensor (gluteus maximus, Glu) and flexor (illiopsoas, Ilps) virtual muscles in NMC-5 during variable-inclination walking (C5), averaged over the six valid participants.
